# Mechanical Stretch Inhibition Sensitizes Proprioceptors to Compressive Stresses

**DOI:** 10.1101/2020.12.30.422571

**Authors:** Ravi Das, Li-Chun Lin, Frederic Català-Castro, Nawaphat Malaiwong, Neus Sanfeliu, Montserrat Porta-de-la-Riva, Aleksandra Pidde, Michael Krieg

**Affiliations:** ICFO, Institut de Ciències Fotòniques, Castelldefels, Spain

## Abstract

A repetitive gait cycle is an archetypical component within the behavioural repertoire of many if not all animals including humans. It originates from mechanical feedback within proprioceptors to adjust the motorprogram during locomotion and thus leads to a periodic orbit in a low dimensional space. Here, we investigate the mechanics, molecules and neurons responsible for proprioception in *Caenorhabditis (C.) elegans* to gain insight into how mechanosensation shapes the orbital trajectory to a well-defined limit cycle. We used genome editing, force spectroscopy and multiscale modeling and found that alternating tension and compression with the spectrin network of a single proprioceptor encodes body posture and informs TRP-4/NOMPC and TWK-16/TREK2 homologs of mechanosensitive ion channels during locomotion. In contrast to a widely accepted model of proprioceptive ‘stretch’ reception, we found that proprioceptors activated under compressive stresses *in vivo* and *in vitro*, and speculate that this property is conserved across function and species.

## Introduction

Many, if not all, motile animals generate forward thrust that is powered by collective cell shape changes due to antagonizing muscle activity. In *C. elegans*, the associated contraction/relaxation cycles are performed with sub-maximal capacity and are driven by four classes of coupled excitatory and inhibitory motor neurons [1], giving rise to the tailward propagating wave that bends the body with a given curvature [2]. The amplitude of these optimal driving patterns is remarkably robust against external and internal perturbations, due in part to mechanosensitive feedback from specialized sensory cells and neurons that signal the mechanical deformation to the central nervous system [3]. Several proprioceptor neurons have been identified in *C. elegans* that become activated upon spontaneous or imposed body bending, such as DVA, PVD, SMD and the motor circuit itself [4–8], all of which express mechanoelectrical transduction (MeT) channels that are likely candidates to read out the mechanical strains and stresses that arise during locomotion [9, 10]. However, we still have little knowledge about the physiologically relevant mechanical stresses and deformations that lead to the activation of mechanosensitive neurons during proprioception or visceral mechanosensation [11]. This is due in part to the complexity of animal anatomy confounded by the coexistence of multiple MeT channels, their differential sensitivity to specific stress tensors [12, 13], and the associated difficulty to record from the mechanosensor in moving specimens. Contrary to intuition, structure-guided modeling revealed that one of the best characterized MeT channels of the TRP family, NOMPC [14, 15], is gated under compressive stresses [16]. In contrast, members of the ubiquitously expressed two-pore potassium (K2P) channels, activate under mechanical tension [17]. Because TRP and K2P channels have opposing roles on neuronal (de)polarization [15], strain selectivity of these ion channels could fine tune neuronal responses in a dynamic environment [18].

Here, we identify that a single neuron, DVA in C elegans, activates upon compression of the spectrin cytoskeleton in a NOMPC/TRP-4 dependent manner, while mechanical tension attenuates neuronal activity and prevents firing through the K2P homolog TWK-16. We speculate that this mechanical interplay is particularly important during body movement to confine neuronal activity within controlled regions when positive and negative stresses coincide in long sensory neurites.

## Results

### *β*-spectrin curbs body bends

We previously identified UNC-70 *β*-spectrin, as a key cytoskeletal component that is under mechanical tension in neurons in *C. elegans* [19]. We noticed that the same mutations that lead to a failure to detect forces during gentle body touch, also increase body curvature during locomotion. Thus, we conjectured that UNC-70 might have roles in proprioception during locomotion. We first recorded short videos of freely moving wildtype (wt) and *unc-70* mutant animals, and applied dimensionality reduction techniques (see Methods and Table S1 for details) to represent the emergent locomotion pattern as a periodic orbit in a low dimensional phase space (Fig. 1A,B and Ref. [20]). The first two modes describe the forward locomotion and form planar, phase locked orbit, with a limit corresponding to the amplitude of the body bends. The third mode has been attributed to turning behavior and deep body bends [20, 21]. These modes, previously termed eigenworms, span a parametric space that effectively describes the stability and dynamics of postural changes within a simple 3D Cartesian coordinate system.

**Figure 1.**
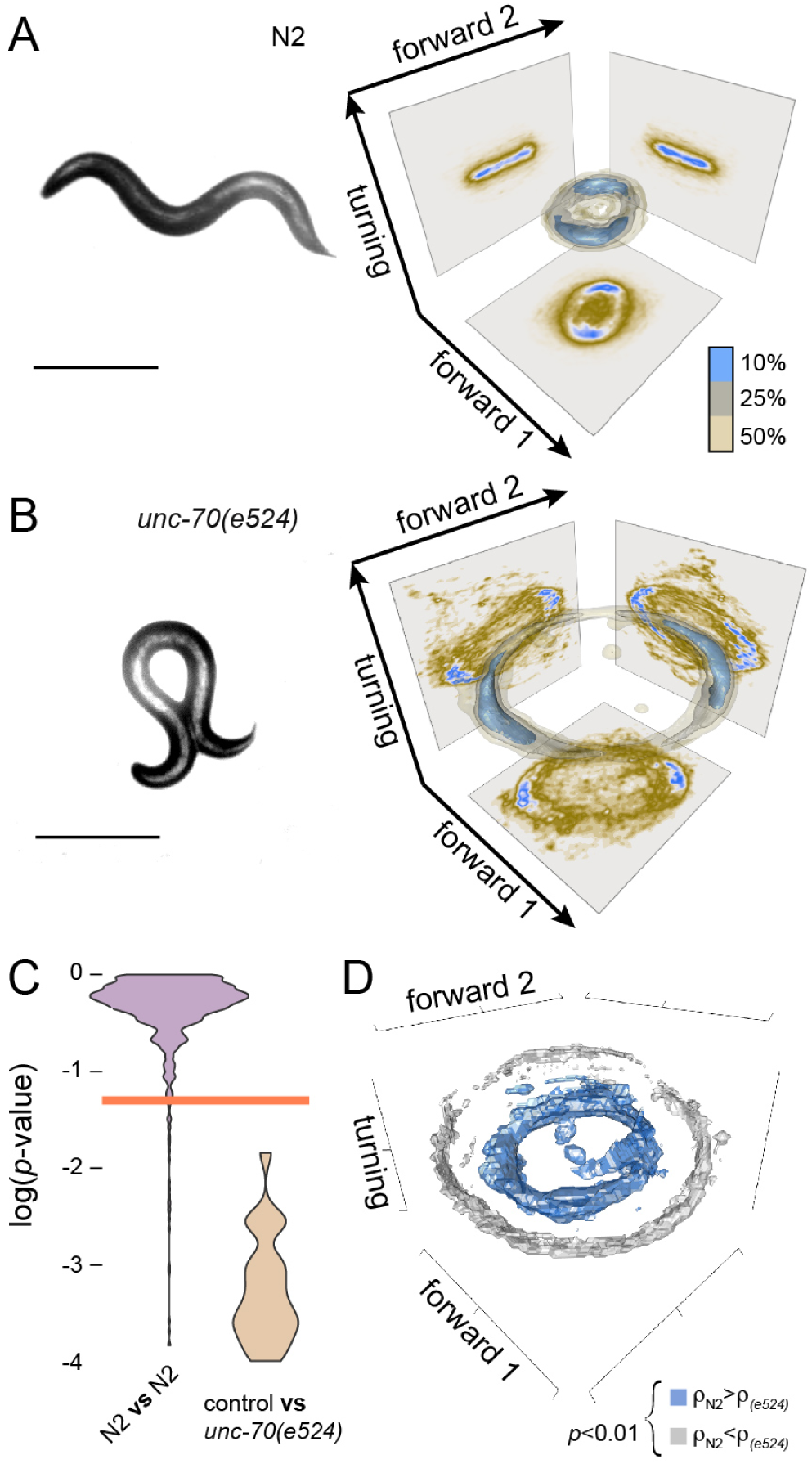
*β*-spectrin constrains the limit cycles during forward locomotion. **A,B** Representative still image of (**A**) wildtype animals and (**B**) *unc-70(e524)* animals and the 3D density estimate for the joint probability distribution (equivalent to a discrete 3D histogram) of the two forward and turning modes in the eigenworm space. Color scale = (brown, low density; blue, high density) **C** Violin plots of the *p*-value distributions for 1000 independent tests of a bootstrapped population estimate of wt 3D probability density function (ctrl) tested against itself or a bootstrapped population estimate of *unc-70(e524)*. Orange line indicates *α*=0.05 level of significance for the hypothesis H_0_ that bootstrapped density functions derived from a and b are equal (see Methods). **D** Color-coded, statistically significant differences in the local probability density functions shown in (A) and (B).

Except for omega-turns, wt animals show little deviations into the third dimension during forward locomotion (Fig. 1A), leading to a planar, toroidal manifold. In contrast, *unc-70(e524)* animals display a much larger orbit in the two forward modes, and significant excursions into the turning mode (Fig. 1B), indicative for severely exaggerated body bends (Fig. 1C,D). With this analysis it becomes apparent that UNC-70 is required to reach and adjust the curvature amplitude during forward locomotion that is optimized for animal locomotion.

### The spectrin network has cell-specific roles during locomotion

Because *β*-spectrin is expressed in many if not all neurons and weakly in body wall muscles [19, 22, 23], a higher bending amplitude could in principle reflect defects in muscle contraction or a loss of neuronal control. We therefore sought to test tissue-specific roles of *β*-spectrin independently and generated a ‘floxed’ *unc-70* allele (Fig. 2A; see Methods for details) that allows for conditional excision upon cell-specific expression of the CRE recombinase in muscles or neurons and confirmed CRE activity using a fluorescent recombination reporter (Ref. [24] and Fig. S1, Table S2). Importantly, neither the CRE lines nor the floxed *unc-70* (Fig. 2B,F, Fig. S2A-C,J) allele had a phenotype on its own when tested separately. Having successfully confirmed CRE activity, we pan-neurally expressed CRE in the floxed *unc-70* background (Fig. 2C, Video S2) and rarely observed a regular orbit in the eigenworm space, indicative for severely uncoordinated locomotion behavior. When we expressed CRE under muscle-specific promoter [25], we did not observe any defect (Fig. 2D) suggesting that *β*-spectrin is functionally restricted to neurons during crawling behavior in adults.

**Figure 2.**
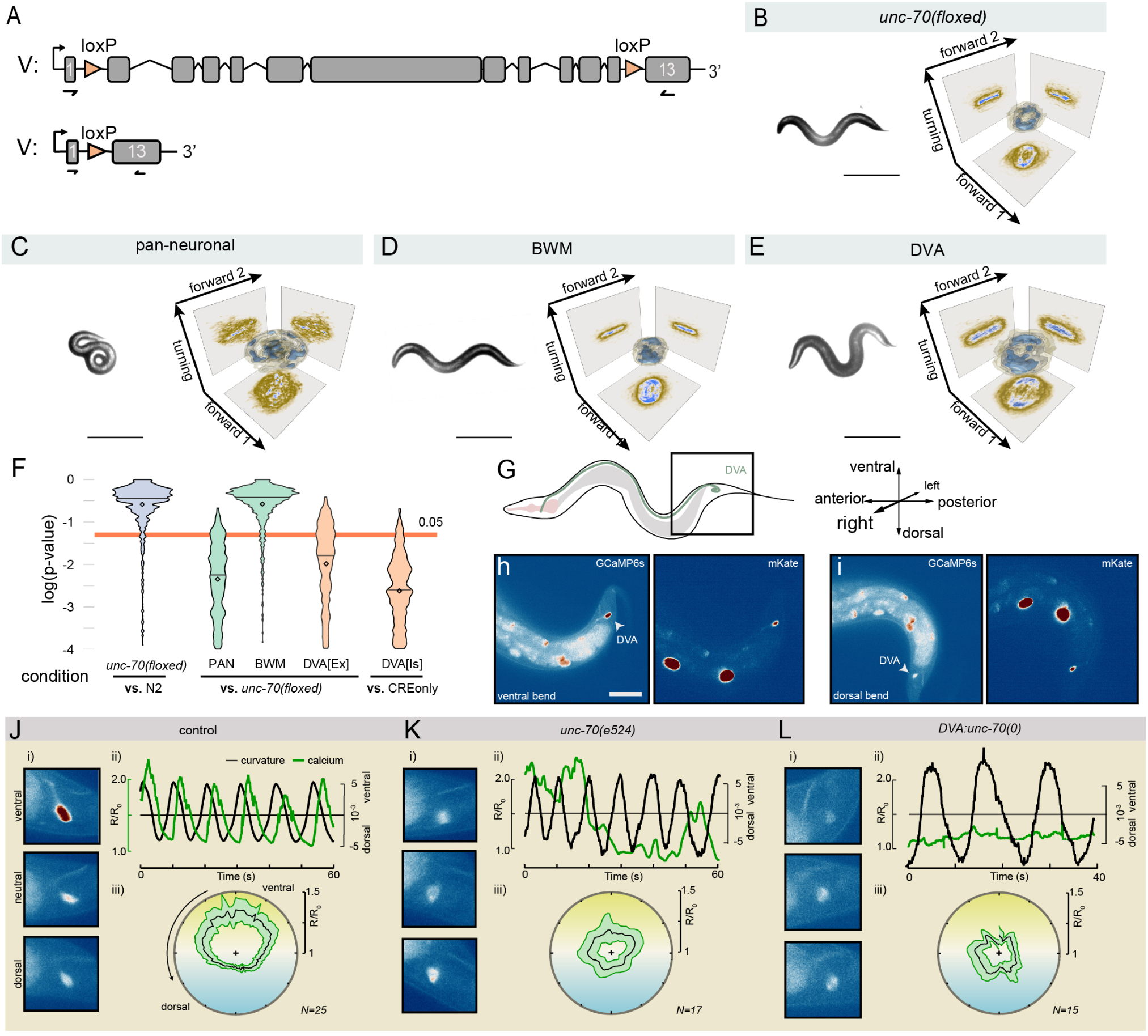
Fig. 2. *β*-spectrin controls body posture specifically in DVA. **A** Genetic strategy for cell-specific deletion of *unc-70* in individual cells and neurons by flanking the *unc-70* genomic fragment with two loxP sites and the resulting genomic scar after recombination. **B-E** Representative still image and the corresponding manifold in the three dimensional eigenworm space for (**B**) *unc-70(loxP*) control animals without CRE expression, (**C**) PAN-neuronal *unc-70* deletion, and *unc-70(loxP)* animals with CRE-expression in (**D**) body wall muscles and (**E**) DVA sensory interneurons. Scale bar = 300 *µ*m. **F** *p*-value distribution for 1000 independent tests for the hypothesis H_0_ that the density function for indicated combinations are equal (see Methods). Orange line indicated *α*=0.05 level of significance. **G** Schematic of an animal with the location of DVA cell body and its axon and the coordinate system of our analysis. For clarity and convenience, we display ventral up to emphasize positive calcium correlation with ventral postures. **H,I** Representative calcium-sensitive (GCaMP6s) and insensitive (mKate) images for (**H**) ventral and (**I**) dorsal body bends. Scale bar=50*µ*m. **J-L** Representative calcium imaging data from moving (**J**) control, (**K**) *unc-70(e524)*, (**L**) DVA::*unc-70(0)* mutant animals expressing GCaMP6s calcium reporter in DVA subjected to ventral and dorsal bends. (i) Video stills of ventral, neutral and dorsal bends. (ii) Time series of the normalized GCaMP/mKate ratio (left axis) and body curvature (right axis). (iii) Average GCaMP/mKate ratio plotted against the phase angle of body curvature. Yellow shading indicates ventral bends (top), blue corresponds to dorsal body posture (bottom).

We then expressed CRE in either A, B or D-type motor neurons or previously proposed proprioceptors SMD, PVD, Touch Receptor Neurons (TRNs) or DVA and confirmed successful recombination (Fig. S1, Table S2). Even though we neither detected a significant locomotion phenotype in A, B and D-type motor neurons (Fig. S2D-F and Video S2) nor in TRNs (Fig. S2G) or PVD (Fig. S2H), *unc-70* deletion in SMD caused subtle defects (Fig. S2I,K) and only the knockout of *unc-70* in DVA (Fig. 2E,F) leads to an abnormally exaggerated body posture (Video 2). This observation motivated us to ask if this DVA-dependent role of UNC-70 in movement was a general property of spectrin fibers or specific to *β*-spectrin. Thus we knocked out SPC-1/*α*-spectrin, using the well established auxin-induced protein degradation (AID) system [26]. We used a previously published SPC-1::AID animal [27] and first verified that the spatially restricted expression of TIR ligase in TRNs leads to a reduction in protein levels upon addition of auxin (Fig. S3A-D). Then, we targeted TIR to DVA and found that neither expression of the TIR ligase (Fig. S3E-H) nor the SPC-1::AID alone (Fig. S3I-L), but only the co-expression of both transgenes in DVA lead to exaggerated body angles, visible as a larger manifold in the eigenworm space (Fig. S3M-P), similar to the CRE-dependent *unc-70* recombination observed before (Fig. 2E). Taken together, our conditional gene ablation strategy unveiled an unexpected cell-specific role, of the otherwise widely expressed spectrin network, in regulating the extent of body bending during locomotion.

### DVA activity correlated with compressive stresses *in vivo*

We next investigated whether changes in body postures influenced neuronal activity in DVA in an *unc-70* dependent manner. We first generated a Ca^2+^ activity reporter using DVA specific expression of Gal4 driving a GCaMP6s effector [28]. Then, we performed live imaging and video tracking to correlate Ca^2+^ activity and body curvature in proximity to the cell body (square in Fig. 2G), and normalized the Ca^2+^ sensitive by a Ca^2+^ insensitive fluorophor (Fig. 2H,I). Even though the low Ca^2+^ signal exhibited occasional spontaneous activity bursts and low-amplitude spontaneous signals in completely restrained animals (Fig. S4A), we observed striking activity changes between the ventral and the dorsal side in animals undergoing full body swings (Video S3, Fig. 2J). To our very surprise, activation did not correlate with dorsal, but strongly with ventral postures, under which the DVA neurites shorten dramatically (Fig. S5A,B).

To understand the role of UNC-70 in this process, we repeated these experiments in the *unc-70(e524)* mutation (Fig. 2K) and in the DVA-specific *unc-70* knockout (Fig. 2L). In both conditions, we did not observe curvature-correlated Ca^2+^ activity, suggesting that *β*-spectrin is needed for neuronal activation as a response to body posture changes and that it acts cell-autonomously within DVA. Rarely, however, signals also appeared during dorsal bends, indicating that UNC-70 directs the preference of neuronal activation during ventral postures.

### *β*-spectrin is under compression during ventral body bends

The finding that DVA activates during ventral bends primed us to investigate how mechanical stresses affect axon shape in moving animals and how the spectrin network contributes to proprioceptive mechanosensitivity. We thus recorded short videos and quantified the local length changes of DVA in each frame as a function of body posture in flexing animals (Fig. S5). Similar to ventral touch receptor neurons (e.g. AVM [19, 29]), DVA locally shortens and elongates up to 40% (Fig. S5A,D) during dorsoventral swings in wildtype animals. In *unc-70(e524)* and DVA-specific *β*-spectrin mutants, however, DVA extended under dorsal body postures but failed to shorten during ventral bends (Fig. S5B-F). Instead, the axon showed a shape change characteristic of a mechanical failure due to compressive stresses, known as buckling instability.

Physical intuition teaches us that an elastic body subjected to bending experiences compression on the convex and extensions on the concave side [30, 31], with a stress *σ* that increases with the distance to the central axis (*d*), elasticity of the worm’s body (*E*) and the curvature assumed (*c*, Fig. 3A). To directly visualize the stresses originating in the body of *C. elegans* in postures that are typical during locomotion, we resorted to a previously characterized FRET tension sensor [19, 32], with predominantly neuronal expression (Fig. SA,B). The inherent mobility of *C. elegans* precluded unrestrained imaging, thus we redesigned a microfluidic device [33] with channels of varying curvatures (Fig. 3B,C) that could trap animals in different postures for 3D imaging (Fig. 3D). In completely straight positions, FRET values were evenly distributed on the ventral and dorsal side of the animals (Fig. 3E). When the same animal was bent inside the channel, the concave side had a higher FRET efficiency than the convex side in a curvature-dependent manner (Fig. 3E,F), indicative for a differential compression and extension of the animal’s body. However, these differences in FRET between the convex and concave side were not observed in a stretch-insensitive sensor that was fused to the N-terminus of the protein, such that it could not be pulled apart and report tension (Fig. 3G,H), or in a constitutive high-FRET and low-FRET construct in which the elastic force sensor domain was replaced with a stiff 5 or 200 aminoacid linker domain (Fig. S6E-H). Likewise, after performing FRET imaging in E2008K spectrin mutant animals (Fig. S6C,D), we failed to detect differences in FRET efficiencies between the compressed (concave) and stretched (convex) side at high curvatures (Fig. 3I,J), indicative for a failure of the mutant *β*-spectrin to sustain compressive and tensile mechanical stresses. Importantly, we conclude that the E2008K mutation does not interfere with formation of the ubiquitous *α*/*β*-spectrin network (see Fig. S6E,F and Ref. [29]), as judged by the *α*-spectrin periodicity in the *β*-spectrin *unc-70(e524)* point mutation. Together, this shows that the spectrin cytoskeleton sustains compressive AND tensile stresses during body bending. We further conclude that changes in body curvature are encoded in the mechanical state of the spectrin network, which conveys mechanical stresses within DVA.

**Figure 3.**
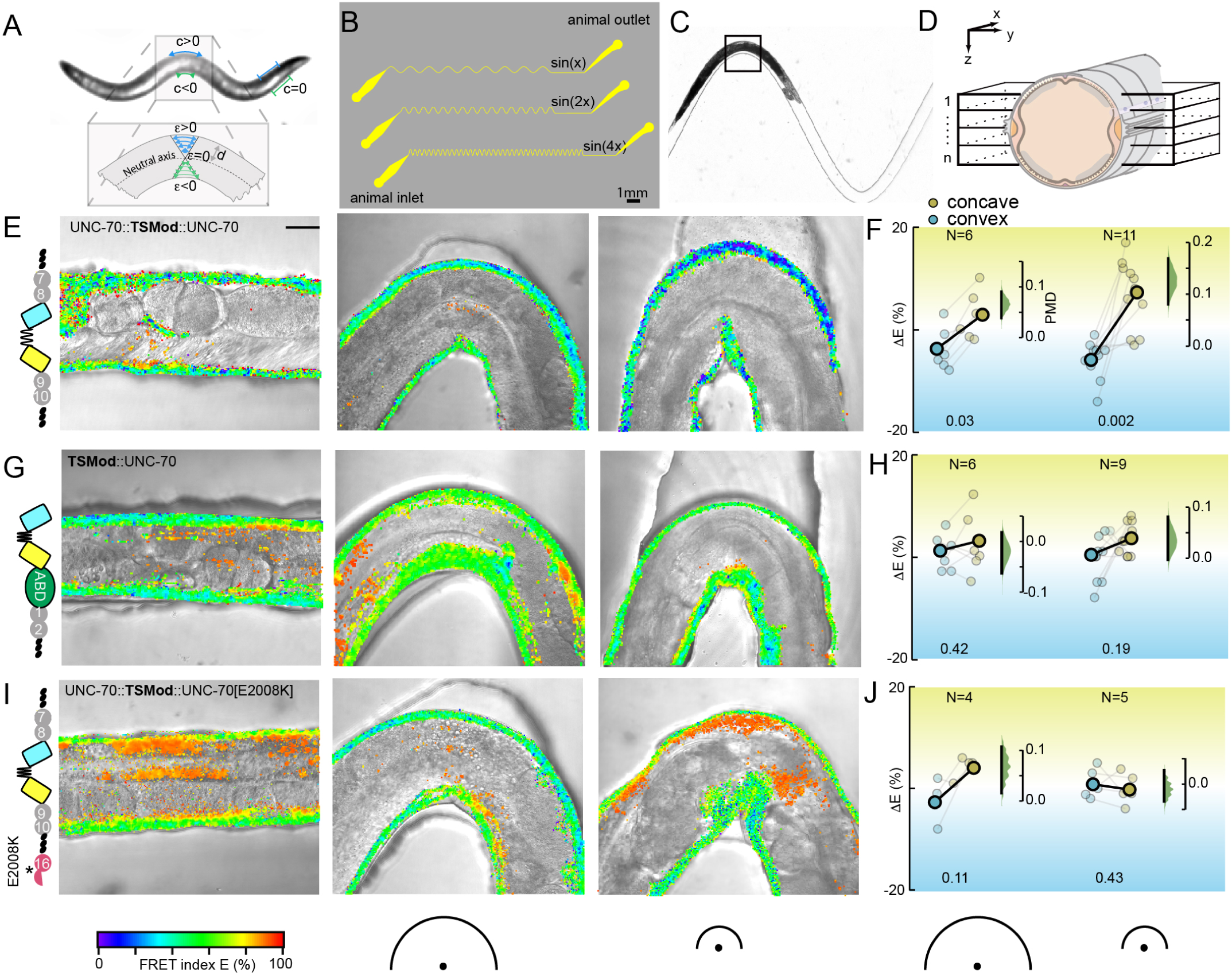
Changes in body mechanics encodes body posture curvature. **A** Schematic of a crawling animal with varying posture along its body and the associated Euler strains. c=curvature; *c*=strain; d=thickness **B** Sketch of the mask layout with varying channel curvatures and periodicities. **C** Brightfield image of an animal inside the sin(2x) channel. **D** 3D representation of the image acquisition procedure. A volume consisting of 10-15 frames encompassing the ventral and dorsal nerve chord was acquired, omitting the lateral nerves. **E,G,I** Schematic and representative and FRET images of collapsed z-stacks for straight and increasingly curved channel of the WFS chip with the FRET index map overlayed ontop of a brightfield image for (**E**) UNC-70::TSMod force sensor with full-length UNC-70 *β*-spectrin bearing TSMod embedded between spectrin repeat 8 and 9, (**G**) force-insensitive FRET control with the TSMod fused to the N-terminus of full length *β*-spectrin and (**I**) UNC-70::TSMod(E2008K) mutant force sensor. Orange pixels indicate autofluorescence from the gut. Colorscale indicating FRET indices; half circles indicating curvatures of the trapping channel, larger circle=smaller curvature. **F,H,J** Quantification of the FRET index difference between the convex and concave side of the body and an uncurved portion of the same animal. Δ*E >* 0 indicates compression, Δ*E >* 0 indicates tension. Each connected pair is derived from one image; bold connected dots indicate mean of the sample. The floating right axis shows the paired mean difference (PMD) as a bootstrap sampling distribution (green) and the 95% confidence interval as a vertical black line. Numbers below the graph indicate the *p*-values for the likelihoods of observing the effect size, if the null hypothesis of zero difference is true.

### UNC-70 genetically interacts with TRP-4 mechanosensitive ion channel

Our notion that DVA-specific mutations of *unc-70* increase body curvature (Fig. 2E) is shared by the function of the mechanosensitive NOMPC homolog TRP-4 in DVA [4], suggesting a functional relation between UNC-70 and TRP-4. Indeed, animals bearing the *trp-4(sy695)*, and occasionally in *trp-4(ok1605)* single mutation (not shown), have significantly different locomotion phenotypes as compared to wt animals (Fig. 4A,D). In order to determine whether or not UNC-70 and TRP-4 function together in determining the body posture during locomotion, we generated animals carrying *trp-4(sy695)* and *unc-70(e524)* double mutations, and *trp-4(sy695)* and the conditional allele with a DVA-specific defect of *unc-70*. In both conditions, the double allele generally did not show a more exaggerated body posture than either single mutation alone (Fig. 4D). Importantly, the distribution and trafficking of endogenously tagged TRP-4 into the distal part of the sensory endings is unaffected by the E2008K spectrin mutation. Expression levels of TRP-4 in DVA remained below background autofluorescence (Fig. 4E) and we restricted our analysis to the cilia of CEP or sensilla of ADE, which locally enrich the protein in sensory endings. Taken together, our data suggests that *unc-70* acts epistatic with *trp-4* to limit body bending amplitude.

**Figure 4.**
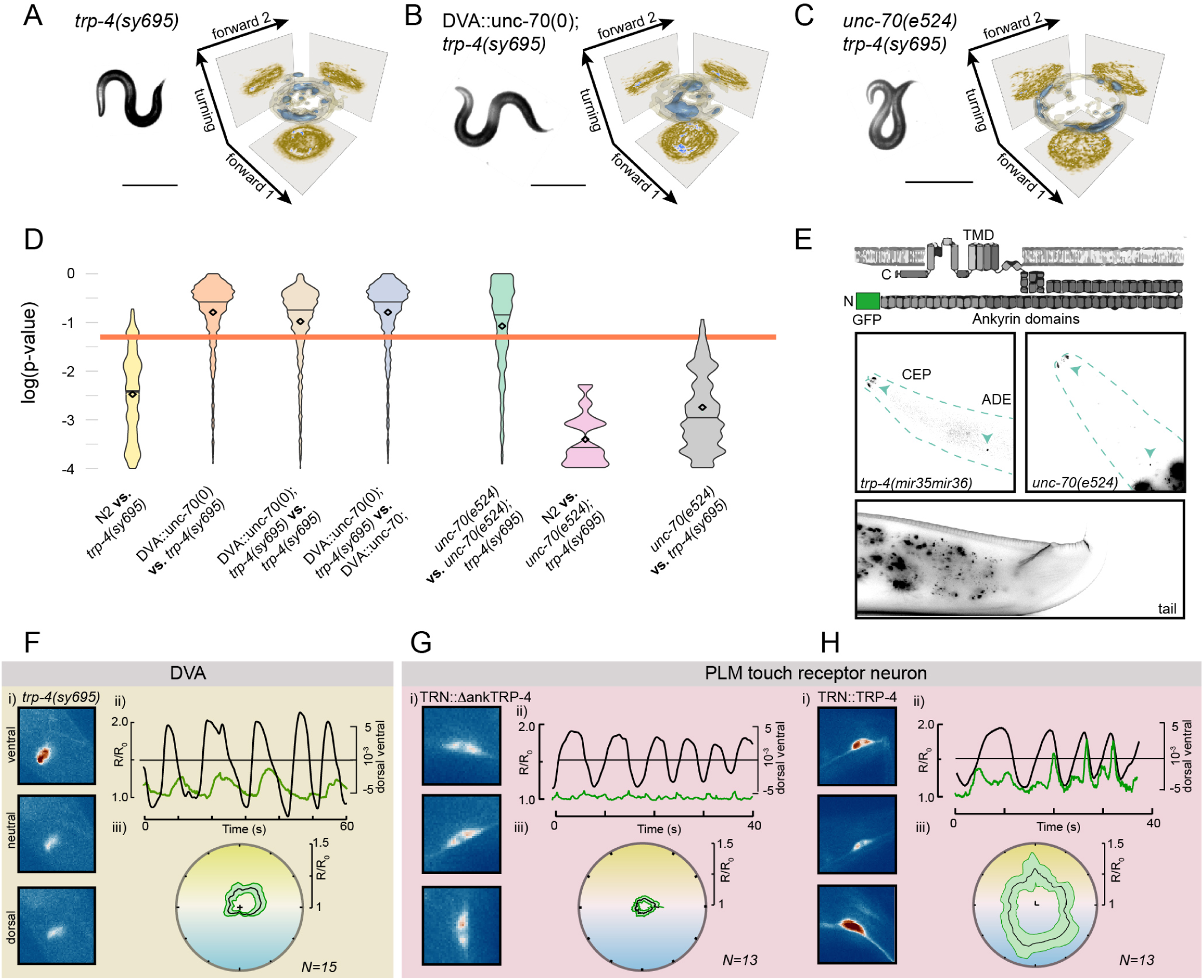
TRP-4 calcium activity peaks under compressive stresses. **A-C** Representative still image and the corresponding manifold in the three dimensional eigenworm analysis for (**A**) *trp-4(sy695)* mutant animals, (**B**) *trp-4(sy695);DVA::unc-70* double mutants (**C**) *trp-4(sy695);unc-70(e524)* double mutation. **D** *p*-value distribution for 1000 independent tests for the hypothesis H_0_ that the density function for indicated combinations are equal (see Methods). Orange line indicated *α*=0.05 level of significance. **E** Schematics of the TRP-4::GFP CRISPR knock-in and representative images of the head and the tail in wildtype and *unc-70(e524)* background. **F-H** Calcium imaging data from moving (**F**) DVA neuron in trp-4(sy695) mutant, (**G**) PLM in an animal expressing N-terminally truncated TRP-4, (**H**) PLM in animals expressing fulllength TRP-4. (i) Video stills of ventral, neutral and dorsal bends. (ii) Time series of the normalized GCaMP/mKate ratio (left axis) and body curvature (right axis). (iii) Average GCaMP/mKate ratio SE plotted against the phase angle of body curvature.

### TRP-4 is not essential to elicit curvature-dependent calcium activity

We next analyzed whether compression induced Ca^2+^ signals depend on the TRP-4/NOMPC ion channel. Indeed, spontaneous activity in immobile *trp-4* mutant animals was strongly reduced, with occasional decreases in Ca^2+^ signals (Fig. S4B). However, when we performed Ca^2+^ imaging in flexing animals, we still observed DVA Ca^2+^ activity during ventral body bends in *trp-4(sy695)* mutant animals, even though, these DVA signals were less modulated by swings in body curvature (Fig. 4F). This analysis shows that TRP-4 expression is dispensable for compression induced Ca^2+^ signal during body bending. We were thus wondering if TRP-4 is sufficient to sensitize otherwise motion-insensitive neurons lacking endogenous TRP-4 expression [34], to activate during dorso-ventral body bends. We hence expressed full-length *trp-4* cDNA and an N-terminally truncated contruct in TRNs, the gentle body touch mechanoreceptors. In wildtype animals and transgenics expressing the truncated isoform, basal Ca^2+^ levels measured in PLM rarely changed and are seemingly uncorrelated with body curvature in slowly moving animals (Fig. S4C,G). However, upon expression of full-length TRP-4 in TRNs, we observed a strong periodic signal in PLM during body bending that correlated well with body bends (Fig. 4H). Interestingly, we also frequently observed increases in PLM activity when the animal bent towards both, the dorsal and ventral sides (Fig. 4H). Taken together, TRP-4 can, in principle, endow curvature sensitivity in heterologous neurons, but is dispensable for curvature induced DVA activity.

### DVA responds to tension and relaxation gradients *in vitro*

We then established a primary culture of *C. elegans* neurons [19, 35] on compliant PDMS surfaces and verified that DVA expressing GCaMP6s robustly activated in response to substrate deformation (Fig. 5A). Clearly, cultured DVA neurons repetitively elicited high Ca^2+^ transients (Fig. 5B,C Video S4), that depended on functional TRP-4 expression and could be partially blocked by non-specific cation channel inhibitor GdCl_3_ (Fig. 5D). Most notable, we frequently found that DVA neuron became active during the force offset (Fig. 5B,C) and buckled during indentation, suggesting that they sensed negative tension gradients or compression. Because PDMS substrate deformation induces compressive and tensile stresses along the axon, we specifically applied positive and negative tension gradients to isolated axons (Fig. 5E,F) and visualized their resultant Ca^2+^ transients using confocal microscopy (Fig. S7A). To do so, we used optically trapped microspheres and extruded single membrane tethers [36] from DVA axons (Fig. 5E) with varying velocity. The resulting tether force increased with the pulling velocity but quickly relaxed to a static value (Fig. 5G) when the movement ceased. Using this approach, we measured local gradients that were 100x higher than the resting membrane tension (2mN/m, Fig. S6B), resulting in a rich behavior in the Ca^2+^ dynamics. Strikingly, Ca^2+^ signals increased preferentially during the relaxation phase of the force-distance cycle directly at the tether neck (Video S5 and Fig. 5F-H), indicating that negative tension gradients, similar to axon compression *in vivo* (Fig. S5A), can induce neuronal activity (Fig. 2J). In *trp-4* mutants, however, we did not observe an increase in Ca^2+^ activity during tension relaxation (Fig. 5I,J). In addition to the Ca^2+^ increases during tension relaxation in wildtype neurons, we also frequently observed a transient reduction in Ca^2+^ signals during tether extrusion especially at higher velocities (Fig. 5F-H), suggesting that the concomitant increase in membrane tension (Fig. 5E-G) suppresses neuronal activity.

**Figure 5.**
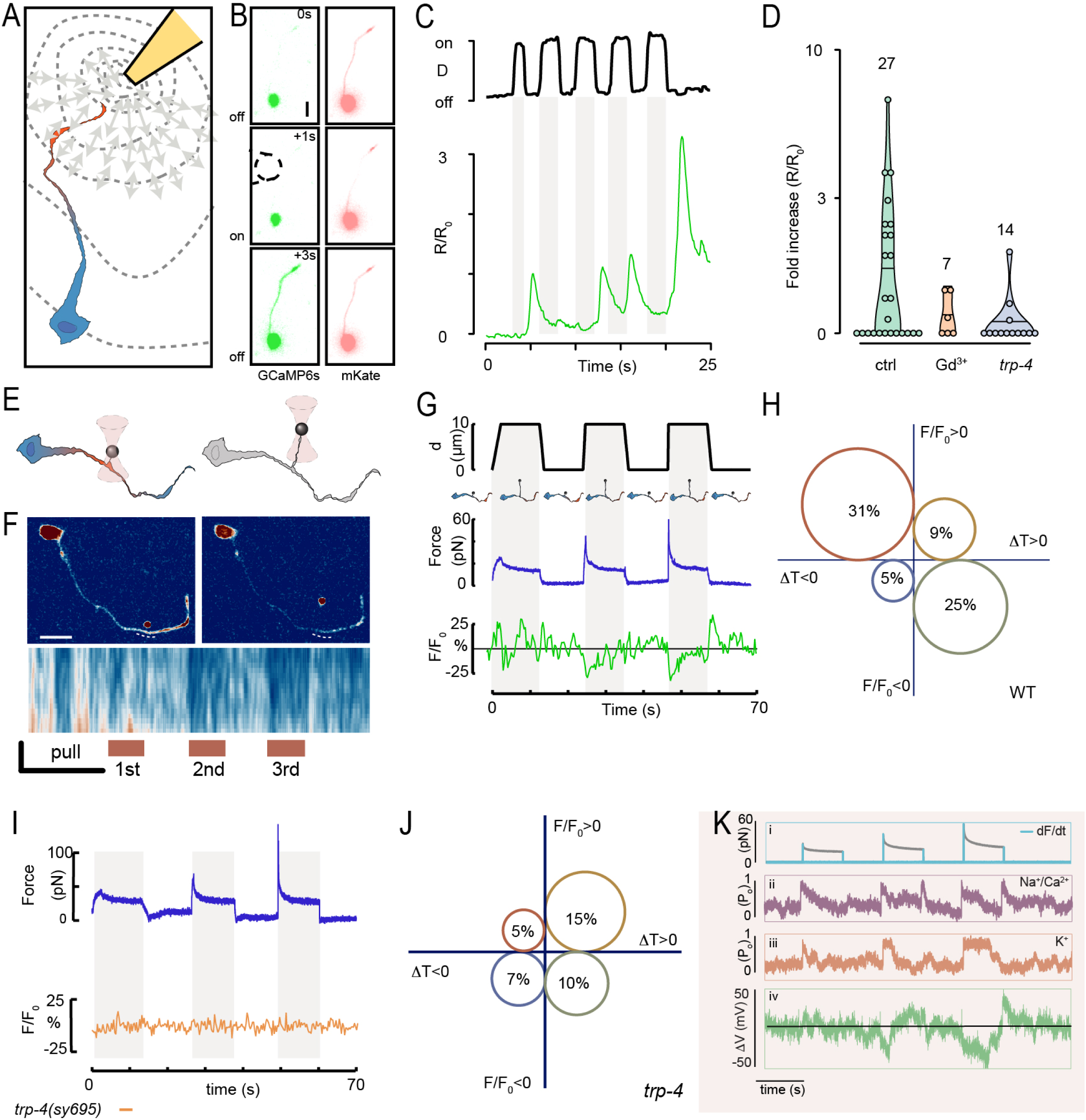
Opposing membrane tension gradients modulate DVA activity *in vitro* through TRP-4 and TWK-16. **A** Schematic of the PDMS stretching experiments. **B** Three successive images of the Ca^2+^-sensitive (left) and Ca^2+^-insensitive dye (right) of an isolated DVA subjected to substrate deformations. **C** PDMS substrate deformation *D* (black line) and GCaMP/mKate ratio (*R*/*R*_0_, green line) plotted against time. **D** Violin plot of the Ca^2+^ activity derived from wt, wt treated with Gd^3+^ and *trp-4(sy695)* mutant DVA neurons. **E** Schematics of the dynamic tether extrusion experiments. **F** Representative still images of the GCaMP signal from an isolated DVA neuron before and during tether extrusion. Dotted line indicates the location of the kymograph displayed below showing the temporal evolution of the Ca^2+^ signal at the tether neck. **G** Representative displacement (*d*), force and normalized Ca^2+^ signal extracted from the experiment in (**F**). **H** Bubble plot of the probability showing that a tether extrusion or relaxation leads to an increase or decrease in GCaMP activity in wt DVA neurons. Δ*T* indicates increases or decreases in measured tension gradient. N=48 cells. **I** Representative force trace and normalized Ca^2+^ transients for *trp-4* (orange) mutant cells. **J** Bubble plot of the probability showing that a tether extrusion or relaxation leads to an increase or decrease in GCaMP activity in DVA neurons mutant for *trp-4*, N=38, mutant cells. **K** *In silico* study of the optical trapping experiment. I) Simulated force (black trace) and temporal tension gradients (light blue). ii,iii) Open probability of the hypothetical Na^+^ and K^+^ channel. iv) Sum of a current through a linear combination of its average open probability x single channel conductance. Note the double peak in the purple trace becomes suppressed due to simultaneous activation of the inhibitory signal, giving rise to the experimentally observed behavior.

How does an increase in membrane tension lead to decrease in calcium signal? This observation, in principle, can be explained in part due to the existence of a stretch activated potassium channel (SAPC) that deactivates the neuron under tension. To understand the observed Ca^2+^ dynamics, we set up a computational model [37], in which positive and negative forces selectively activate hyperpolarizing and depolarizing ion channels, respectively. We illustrate this scenario with two hypothetical, mechanosensitive ion channels acting as primarily sodium/calcium conductive (depo-larizing) and a potassium conductive (hyperpolarizing) and model channel gating under force as a thermally driven escape over a potential barrier [38] (Fig. S7E, for details see Methods). Consistent with previous results derived from electrophysiological recording of TRP-4 expressing CEP [39], we modeled sodium activity at the force onset and offset by sensitizing it to the loading rate (Fig. 5Kii) while K^+^ channel displayed activity only during force onset (Fig. 5Kiii). With realistic ion channel parameters taken from the literature (Methods) and assuming a constant input resistance, the open probability of the K^+^ channel was able to completely suppress the Ca^2+^ channel induced neuron activity at the force onset but not when the force was released. Even though our calculation does not include the complex regulation and spontaneous Ca^2+^ dynamics that likely take place inside a cell, our reductionist model is in good match to the experimentally determined DVA activity (Fig. 5G), such as a transient reduction during extension and transient rise due to relaxation. We next sought to test this prediction.

### The K2P homolog TWK-16 suppresses Ca^2+^ activity during dorsal posture

The results of our simple kinetic model predicts that the Ca^2+^ signals at the force onset becomes unmasked in absence of inhibitory, potassium activity. DVA is one of the few neurons that expresses the mechanically regulated leak potassium channel TWK-16 [40, 41], a TREK2 homolog of mechanosensitive, two-pore K^+^ channels [42] in *C. elegans*. We thus repeated the dynamic tether extrusion experiment in a *twk-16* null mutant background (Fig. 6A,B) and measured the concomitant change in GCaMP intensity. Consistent with our hypothesis, we did not observe a Ca^2+^ decrease during tether pull-out, indicating that mechanical de-activation at high membrane tension gradients is *twk-16* dependent. Instead, we frequently observed a Ca^2+^ increase at the onset of the tether extrusion (Fig. 6A,B), as if TWK-16 functions in suppressing tension-induced activity normally observed in TRP-4 expressing neurons [39]. The notion of stretch-induced suppression of Ca^2+^ activity through TWK-16 in isolated DVA neurons, raises the question of whether or not TWK-16 is a functional component during proprioception. To answer this, we first repeated the Ca^2+^ imaging of DVA in moving *twk-16* mutant animals. Compared to wt animals, we found that the deletion of *twk-16* caused a subtle increase in Ca^2+^ activity in DVA during spontaneous body bending (Fig. 6C) with an overall unchanged ventral preference. Nevertheless, we frequently observed GCaMP intensity changes during dorsal posture and even during both, dorsal and ventral posture (Fig. 6C), a signature we never observed in DVA of wt animals but occasionally in *unc-70* mutants (Fig. 2L) and ectopic expression of TRP-4 in TRNs (Fig. 4H). Together, this suggests that TWK-16 is able to suppress Ca^2+^ transient at dorsal posture, when DVA experiences mechanical tension, similar to how tension suppresses calcium activity in our *in-vitro* experiment (Fig. 6A).

**Figure 6.**
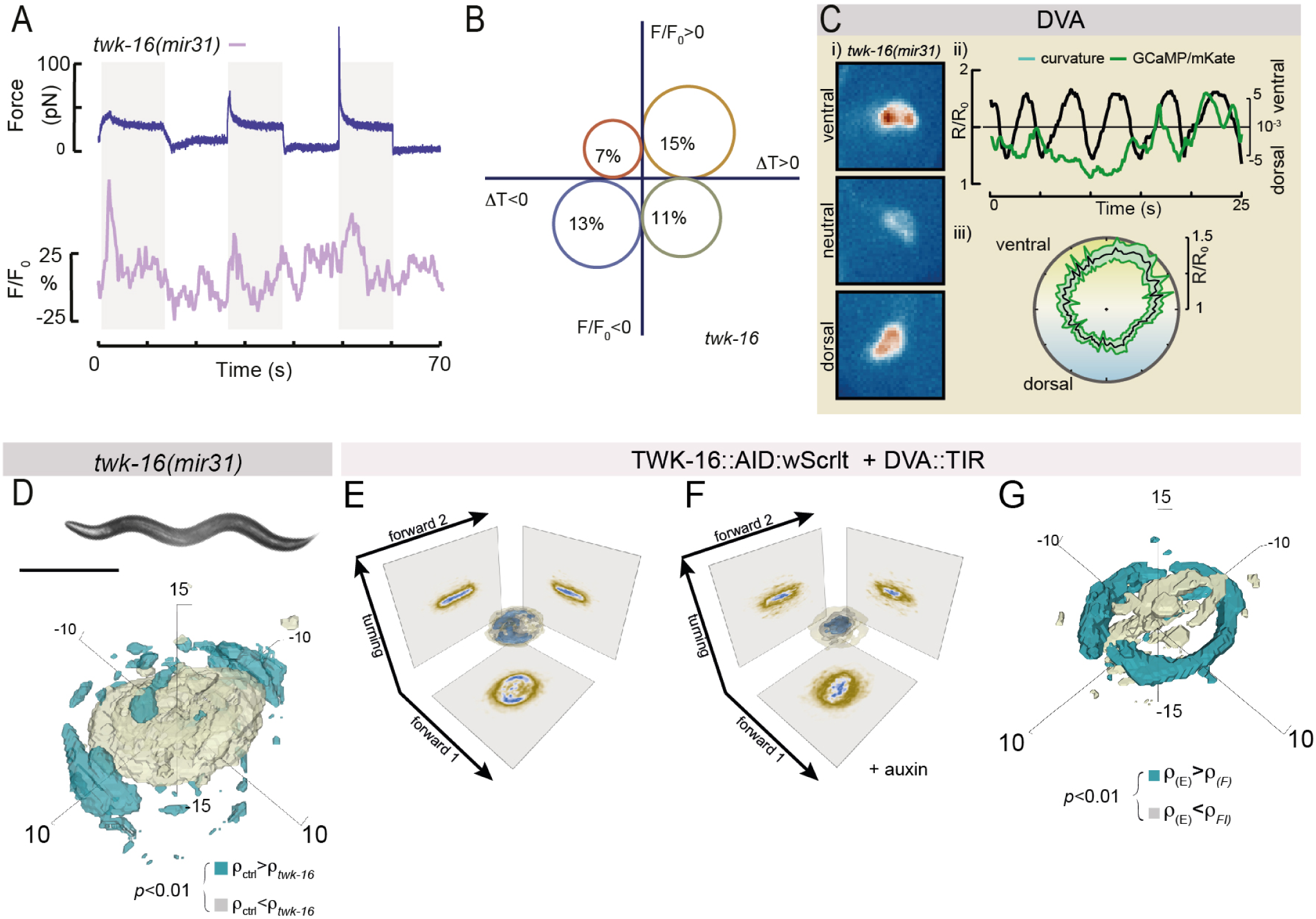
TWK-16 suppresses bending excursions through modulation of DVA activity. **A** Representative force trace and normalized Ca^2+^ transients for *twk-16* (purple) mutant cells. **B** Bubble plot of the probability showing that a tether extrusion or relaxation leads to an increase or decrease in GCaMP activity in DVA neurons mutant for *twk-16*, N=31, mutant cells. **C** DVA Ca^2+^ data of *twk-16(mir31)* animal lacking functional TWK-16. (i) Video stills of DVA cell body during ventral, neutral and dorsal bends. (ii) Time series of the normalized GCaMP/mKate ratio (left axis, green) and body curvature (right axis, black). (iii) Average GCaMP/mKate ratio ± SE plotted against the phase angle of body curvature. **D** Representative image of a *twk-16(mir31)* animal and the color-coded, statistically significant differences in the local probability density functions between N2 and twk-16. Scale bar = 300*µ*m. **E,F** 3D kernel density function (histogram) in the three dimensional eigenworm space for TWK-16::AID::wScarlet animals expressing TIR in DVA in (**E**) absence and (**F**) presence of auxin. **G** Color-coded, statistically significant differences in the local probability density functions between (**E**) and (**F**). Blue voxels indicate higher local density for untreated, beige voxels indicate higher density for auxin-treated animals on the *α*=0.01 level.

We then asked if TWK-16 expression in DVA is required for animal locomotion. In contrast to *trp-4* animals, *twk-16* null mutants, we observed a subtle, but significant difference between the wt and mutant in 3D densities in the eigenworm space, as an indicator for a smaller body posture (Fig. 6D, S8A-C). TWK-16 is also expressed in other neurons, including AVK that counteracts DVA activity [21]. First, we inserted a AID:wScarlet tag [26] at the TWK-16 C-terminus, verify expression in DVA axon and cell body (Fig. S8D), and confirmed that the fusion protein did not cause any locomotion defects in absence and presence of auxin (Fig. S8E-G). Then, we expressed the previously characterized TIR ligase (Fig. S3I-L) together with the TWK-16::AID and repeated the behavior experiment. Even though we already noticed an auxin-independent effect [43] on body bending amplitude (Fig 6E) the addition of 1mM auxin caused a markable reduction of body bending amplitude compared to wt and a clearly distinguishable eigenworm representation (Fig. 6D,E; S8H-J; Video S6). Together, TWK-16 inactivates DVA under tension and counteracts TRP-4 depolarization with DVA-specific function in locomotion.

### Compression-induced activity adjusts motor output in a neuronal network model

The known wiring diagram of *C. elegans* offers the opportunity to ask how the observed neuronal activity changes are processed on the nervous systems level. To interrogate how the compressive proprioception of DVA adjusts the motorcircuit (Fig. 7A) and affects *C. elegans* locomotion, we adapted a well-established neuromechanical model [44] that was previously deployed to simulate the effect of stretch-sensitive feedback embedded directly into the motorneurons. In our modification, we consider a single compression sensitive current during ventral bends, matching our notion that DVA is excited during ventral curvatures. In agreement with the connectivity of the *C. elegans* network, DVA directly informs the motor neurons on the dorsal side. Indeed, in this scenario, the simulation of compressive-sensitivity during ventral bends, but not stretch-sensitivity during dorsal bends produced a crawling pattern seemingly similar to experimental results (Fig. 7B). With the aim to understand how the compression-current sensitivity influences crawling pattern, we reduced the responsiveness to compression, compatible to the *unc-70* and *trp-4* mutations in the experiments (Fig. 4). Strikingly, the simulation recapitulated the crawling behavior observed in the mutant conditions, visible as an expansion of the manifold in the eigenworm state space (Fig. 7C, Video S7).

**Figure 7.**
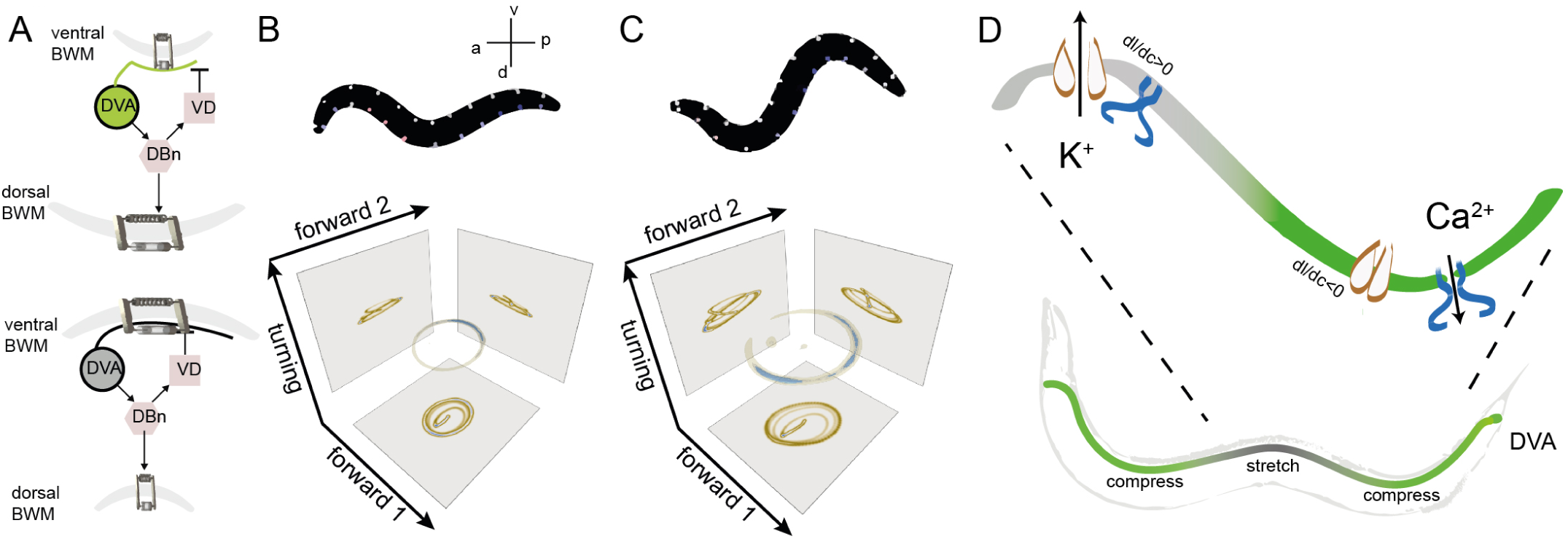
Compression in DVA provides proprioceptive feedback to the motorcircuit. **A** Schematic of the model formulation. A passive mechanical body is modeled as a lightly damped elastic beam (passive Kelvin-Voigt material) [44] in parallel with the muscles, as an ‘active’ Kelvin-Voigt material with stiffness governed by the state variable of the motorcircuit (through neuronal input). DVA senses compression during ventral bends and informs the DB neurons, which in turn activate dorsal muscle contraction and at the same time connect to inhibitory VD motor neurons, resulting in the relaxation of the ventral muscles. Under dorsal bends, DVA activity is suppressed (lower panel). **B,C** A representative snapshot from the simulation of (**B**) ‘wildtype’ and (**C**) ‘mutant’ with reduced sensitivity to compressive stresses and the corresponding manifold in the eigenworm state space. Dorsoventral circles depict motorneurons and their stretch receptor input. Blue=off, red=on. Ventral motorneurons are white, as they do not receive stretch receptor input. **D** Illustration of the mechanical scenario under which neuronal tension hyperpolarizes (activates TWK-16/TREK potassium channels, orange) and compression depolarizes (TRP-4/NOMPC, blue) neuronal segments. Hyperpolarized domains thus limit the extend and spread of the signal along the length thus leading to effective compartmentalization.

## Discussion

Proprioceptive and visceral mechanosensation are vitally important processes ensuring body homeostasis and organ function. Here, we genetically, mechanically and computationally deciphered the molecular underpinnings that determine the shape of the limit cycle attractor during *C. elegans* locomotion. We used a well characterized mechanical stress sensor [19, 45] in combination with neuronal activity indicators and revealed that DVA activates under compressive stresses in an UNC-70 *β*-spectrin dependent manner. This apparent discrepancy in light of the classical view of mechanical stretch reception is not a sole feature of DVA but has previously been observed in other *C. elegans* proprioceptors such as SMD [6], *Drosophila melanogaster* multidendritic class III proprioceptors [46] which preferentially activate in a direction-selective manner when the contracting segment gets compressed [46], and mechanosensory neurons regulating thirst in the mammalian brain [47]. Latter is directly sensed by volume changes through a specific TRPV1-microtubule interaction when neurons lose water the membrane-bound ion channels push against the highly interweaved microtubule cytoskeleton, thus activating the neurons [47]. It seems plausible that this ‘push-activation’ model also accounts for the activation of the pentascolopedial chordotonal organs in the fruitfly, which have been shown to activate under compressive stresses during locomotion [48] in a NOMPC dependent manner. Intriguingly, the TRP family member NOMPC also interacts with a specialized microtubule cytoskeleton [49, 50] which conveys mechanical stress to the gating pore through a conserved ankyrin spring. Whether or not tension [51] or compression [16, 52] of the ankyrin repeats leads to mechanical gating of the pore, remains to be determined, nonetheless, the application of membrane tension was not sufficient to activate mammalian TRP channel homologs [53]. Likewise, our results from the truncated TRP-4 lacking ankyrin domain (Fig. 4G), suggests a membrane-independent activation mechanisms. TRP and TMC channels are not the only ones implicated in compression sensing. Recent work demonstrated that a normal force of 50pN is sufficient to cause conformational change in PIEZO1 [54], providing a plausible mechanism for compressive mechanosensitivity [55, 56]. Our work showing compressive mechanosensitivity *in-vitro* and *in-vivo* thus motivates us to rethink the operation of ‘stretch’ sensitive mechanoreceptors. Future work needs to directly address if TRP-4/NOMPC ion channels are gated by compression of their ankyrin spring.

What interferes with DVA mechanosensing in the spectrin mutation? Our genetic analysis suggests that the spectrin network and TRP-4 act in the same process, e.g. during force transmission. We performed a co-immunoprecipitation of TRP-4 and UNC-70 as well as SPC-1 from CHO cells, but could not detect a direct biochemical interaction in this assay (not shown). Hence, we propose that indirect force transmission takes place involving stress propagation through local and global axon mechanics. We showed that DVA axons in *unc-70* mutations undergo buckling instabilities (Fig. S5), which could, similar to events in TRNs, cause a dramatic reduction in the number and length of microtubules [29], consistent with curvature-induced fracture during buckling [57]. This disintegration and disassembly of the load bearing microtubule filaments during cyclic strains conforms to fluidization of the cytoskeleton [58], which constitutes a plausible mechanism of how a single point mutation in spectrin could interfere with efficient force transmission to the ion channel: the loss of tension sets the origin of the fluidization with vanishing stiffness [59] and thus a failure to built-up and convert compressive stresses from the cytoskeleton into ion channel opening.

If a single mechanosensory neurite integrates information from mechanical compression and extension, that also need to be processed locally, mechanisms need to be in place to restrict spread of depolarization. DVA has a single neurite that spans the entire length of the animal’s body, such that tensile and compressive stresses coexist during dorso-ventral body bends. Under the notion of a high input resistance inherent to *C. elegans* neurons [60], a small input current during compression could in principle delocalize and depolarize the entire axon, hence disrupting the proprioceptive coordination along the length of the animal. We propose that stretch activates TREK-2/TWK-16 and hence suppresses or at least dampen the NOMPC/TRP-4-related transduction current to achieve compartmentalized alternating ‘active zones’ that correspond to the compressed state of the neurite (Fig. 7D). Such a mechanism would not only facilitate gain control during mechanical signaling subjected to stochastic ion channel noise but also allows to sample the full dynamic range of proprioceptor activity.

At least two alternative processes could plausibly explain the observed Calcium increase during ventral bending: First, TRP-4 and TWK-16 could both be activated under mechanical tension during neurite extension on dorsal bends, but close with different kinetics. Under the assumption that TRP-4 closing kinetics are slower than TWK-16, they would remain open for a longer time and an apparent ventral activity is obtained. Our in vitro data showing that Ca^2+^ activity increased during tension relaxation can be interpreted as an argument against this scenario. Second, a single synapse from the ventral motor neurons to DVA has been described [61]. This notion allows us to hypothesize that the ventral Ca^2+^ increases in DVA could be due to corollary discharge from VB motor neurons that provide a copy of the motor state to the central nervous system [62]. Our data showing that DVA-specific mutations in unc-70 lead to decorrelated Ca^2+^ dynamics contradicts this interpretation. However, whole animal Ca^2+^ imaging experiments are needed to further investigate and completely disprove this scenario.

In summary, our data revealed that compressive and tensile stresses in the spectrin network modulate two opposing, excitatory and inhibitory ion channels of DVA proprioceptors, TRP-4 and TWK-16 respectively, that are critical to confine the full, deep modulation of Ca^2+^ activity in moving animals. We suggest that this may be a general mechanism by which long mechanosensory dendrites achieve local computation through mechanical compartmentalization. Future experiments will need to directly address how this mechanism transcends the animals kingdom and human gait adaptation.

## Acknowledgments

We would like to thank the NMSB and SLN labs for discussions and suggestion throughout the work and their use of microscopes and the ICFO biolab and nanofabrication facility for support with animal maintenance and SU8 lithography. We thank Manuel Zimmer, Miriam Goodman, Erin Cram, Sohei Mitani and Sander van der Heuvel for donating animals, Shawn Lockery for microfluidic devices, Martin Harterink and Liu He for help with protein biochemistry, Miriam Goodman and Martin Harterink for critical comments on the manuscript and the CGC (National Institutes of Health - Office of Research Infrastructure Programs (P40 OD010440)).

## Author contributions

RD: animal husbandry, molecular biology and CRISPR, CRE recombination and locomotion, analysis and writing. LL: building animal behavior tracker, writing software for Calcium and locomotion analysis, first draft. FCC: microscopy, optical trapping, tissue culture, analysis, code writing, first draft. NM: animal husbandry, molecular biology, behavior assay and calcium imaging, first draft. NS: animal husbandry, molecular biology. MPR: animal husbandry, molecular biology. AP: Neuromechanical modeling. MK: Concept and acquisition of funding, analysis, programming and writing.

## Code and Material availability

All reagents produced are freely available upon reasonable request to the corresponding author. Some strains will be deposited to the CGC. Scripts developed supporting the analysis can be accessed under Gitlab::NMSB and all relevant behavioral recordings will be deposited to Zenodo upon publication.

## Supplementary Material

### 1 Materials & methods

#### 1.1 Soft lithography and PDMS replica molding

SU-8 soft lithography and PDMS replica molding have been done as described before [63]. In short, the fabrication of the molds was undertaken in-house as a single layer process using standard SU-8 photolithography techniques. We first applied a 5 *µ*m thick adhesion layer to reduce lift-off of the patterned structure during device fabrication. Piranha cleaned 4 inch wafers were used to create an adhesion layer using SU-8 50 before photo patterning in SU-8 2000. After fabrication, molds were vapor-phase silanized in chlorotrimethylsilane to prevent adhesion of the PDMS to the substrate. A 10:1 mixture of Sylgard 184 prepolymer/curing agent was degassed (≈30min in vacuum desiccator) and poured onto the silanized molds. After settling, the PDMS/wafer were baked at 80°C for two hours. Devices were then cut using a scalpel, lifted off and punched with a biopsy punch (1mm or 0.75mm). The procedure of animal insertion into the trapping channel has been described in detail elsewhere [63]. In brief, to load individual in the chip, one young adult worm transgenic for TSMod in UNC-70 and its derivatives (GN517, GN519, GN600, MSB233; Ref. [19]) were picked from an NGM plate containing OP50 bacteria and transferred to a 5 *µ*l droplet of 30% Optiprep (to reduce scattering from PDMS due to refractive index mismatches between the animal and the surrounding [65, 66]) placed onto a hydrophobic substrate (25cm^2^ Parafilm) to swim for 30 sec and rid themselves from bacteria. Then, using a stereo dissecting scope at 60x total magnification (Leica S80), the animals were aspirated into a 23 gauge metal tube (Phymep) connected to a 5 ml syringe (VWR) with a PE tube (Phymep, BTPE-50, 0.58×0.97mm) pre-filled with 30% Optiprep buffer. The loading tube was inserted in the inlet port of the device, while a gentle pressure onto the plunger of the syringe released the animals into waveform sampler. Channel with a thickness of 60 *µ*m were used to firmly hold animals immobile during the 3-channel FRET imaging procedure without stretching or confining them visibly. After each image, the animals were pushed for a distance of *π* into the device to probe the opposite curvature at the same body coordinate.

#### 1.2 Animal maintenance

Animals were maintained using standard protocols [67, 68] and grown at 20°C, unless indicated otherwise. For Gal4 expressing transgenes, animals were maintained and raised at 25°C to ensure consistent expression [28].

#### 1.3 FRET imaging

Three channel FRET imaging of animals transgenic for unc-70(TSMod), unc-70E2008K(TSMod), no-force and high-FRET controls was carried out as described previously [19]. Importantly, the 2000 basepair *unc-70* promotor fragment drives our *unc-70* cDNA predominantly in neurons, with little expression in muscles and no expression in the hypodermis. In contrast, visualizing *unc-70* expression from the endogenous locus with a c-terminal slowtimer fluorescent protein [70] showed significantly expression in the hypodermis and muscles (not shown). In short, animals were either immobilized on agar pads [71] or in microfluidic chips with varying curvature as described above. Imaging was carried out on a Leica DMI6000 SP5 confocal microscope through a 63x/1.4 NA oil immersion lens. Three images were acquired, the direct donor (mTFP2) excitation and emission, donor excitation and acceptor emission and the direct acceptor (mVenus) excitation and acceptor emission. mTFP2 was excited using the 458nm, while mVenus was excited with the 514nm line of an Argon ion laser at 80% and 11% transmission respectively (with 25% of its full available power). The fluorescent light was collected using two hybrid GaAs avalanche photodiodes with 100% gain through an acusto-optical beam splitter with the donor emission window set between 465-500nm and the acceptor emission window set between 520-570nm. Linearity of the detector and the ratio 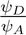 (ratio of collection efficiency of the two APD) was determined experimentally by imaging a homogenous fluorescein solution with increasing gain at constant laser power. Each image was acquired as an average of 4 frames acquired at a 400Hz line-rate with 512×512 pixels at a digital zoom 3 (for Figure S6) or zoom 1-1.5 (for Figure 3) in the microfluidic chip. For each imaging session, the donor->FRET bleedthrough and acceptor cross-excitation by the donor laser was determined using a sample with mTFP (ARM101 or MSB60; [72]) or mVenus (GN498) expression only, respectively, [19]. The bleedthrough and crosstalk factor were calculated as described and assumed to be constant across all intensities and images (for a given set of parameters). The raw FRET intensity was then corrected according to

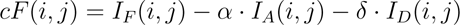

and the FRET efficiency calculated using

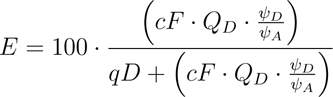

For whole animal images in the WFS, collapsed maximum intensity projections from 15 images of a 3D stack undersampled with ≈2 *µ*m interplane distance were background subtracted with a constant value determined in an ROI that did not contain information from the animal. Before FRET calculation, the individual images were binned to 256×256 pixels to increase signal/noise ratio. For final display, the FRET map was overlayed ontop of the corresponding brightfield image and displayed in Figure 3. In contrast, individual neurons were traced using a Gaussian fit to the intensity profile and processed as described [19].

#### 1.4 Neuron morphology imaging and analysis

Animals expressing mKate in DVA were embedded in the low percentage agar (1-2%), which allowed animals to undergo dorso-ventral body bends without moving out of field of view. Imaging was performed on a Leica DMi8 through a 40x/1.1 water immersion lens using the 575 nm line of a Lumencor SpectraX LED lightsource. Fluorescence was collected through a 641/75nm emission bandpass filter (Semrock, FF02-641/75-25) and recorded using a Hamamatsu Orca Flash 4 V3 for minutes at 10Hz with an exposure time of 50ms. Out-of-focus frames were discarded and the resulting videos were post-processed as previously described [19].

##### Locomotion behavior analysis

###### Animal tracking platform and data acquisition

Animals were synchronized using standard alkaline hypochlorite (bleaching solution) treatment method. Arrested L1 were seeded onto OP50 NGM plates and incubated at 20°C. Without a lack of generality and to facilitate automated tracking and post-processing, we recorded videos of the locomoting animals without food (OP50). To ensure that the absence of food did not alter the outcome of the experiment, we sampled the mutant genotypes and wt on food 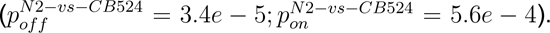 Each video containing a single animal was 1-2 minutes in length and was taken at 25 frames per second (fps). In total, ≈ 1 million frames were collected and processed (Table S1). Videos of the young-adult animals were taken within 5 min after picking using a home-built animal behavior tracking platform. The custom build macroscope is composed of an sCMOS camera (IDS UI-3080 CP Rev. 2) coupled to a Navitar 6.5x zoom lens at 2x zoom. A closed loop stage tracking algorithm, implemented in C#, kept the animal in the field of view for the duration of the recording, or was manually followed with a motorized xz-scanning stage (102×102mm travel, Standa), trans-illuminated with a diffused white light LED array.

###### Auxin experiment and quantification of the knockdown

Auxin plates were prepared by adding 100 mM stock of 1-Naphthaleneacetic acid (Auxin, Sigma Aldrich 317918) dissolved in 100% ethanol to cooled NGM right before pouring the plates at a final concentration of 1mM auxin. Once solidified the auxin containing plates or only ethanol (1%) containing control plates were seeded with concentrated 10X OP50. Young-adult animals containing TIR plasmid injected at 10 ng/*µ*l were transferred to the auxin or control plates and incubated at RT for 2 hours prior to the video acquisition of their locomotion behavior.

###### Data analysis using eigenworms

Imaging processing was performed using custom MATLAB scripts based on Ref. [73] using the following methods:

1. All video frames were converted to grayscale images showing only the worm itself with a white background, which was achieved by background subtraction and two thresholding procedures.
  a. Background subtraction Each video was divided into several equal blocks. Then a background image was created by taking the brightest pixel from the images, which was used to remove the background from the frames from that block.
  b. First thresholding After the background subtraction, the first threshold value was applied to the images that resulted in removal of most of the dark non-worm pixels.
  c. Second thresholding Subsequently, an adaptive thresholding, also known as Bradley method [74], was applied to remove the remaining small clutters, specially close to the worms.
2. Fast tracking pipeline, adapted from Ref. [73], was applied. In brief, all frames were passed through the fast tracking pipeline, however only simple (non-self-overlapping posture) frames were successfully processed, but concurrently labeling the unprocessed frames as ‘crossed’ (self-overlapping posture) frames which required further processing. The results from the tracking pipeline for the simple frames include the head/tail position, backbone, thickness and modes (*a*_1_, *a*_2_, *a*_3_).
3. Crossed frames tracking
  a. Manual tracing First, ‘crossed’ frames were divided into a number of blocks where each block consists of continuous ‘crossed’ frames. Then, each ‘crossed’ block was down sampled at 5 Hz to perform manual annotation of the backbone.
  b. Comparative reconstruction For comparative reconstruction, we used either the first manually traced or the last image of the simple block (if any) to align with the first image of the crossed block using cross correlation, and to calculate the difference between the two images. Then, we applied the image thinning process to find the backbone of this difference. This backbone was then connected to the rest of the worm’s backbone for that corresponding image. The comparative reconstruction step was repeated four times until the next manually traced frame. Now, the new manually traced frame served as the reference for next four frames and the process continued until the end of the crossed block. After the comparative reconstruction we only got backbone of the worms, thus we used the backbone to calculate the required modes (*a*_1_, *a*_2_, *a*_3_).

##### Calcium imaging and analysis

###### Imaging

Young adults animals were mounted onto 1-1.5% of agar pad with 3-5 µL of latex beads (Polybeads, 0.2 µm, PolySciences) to facilitate body movement and stabilize the position of field of view [71]. Calcium imaging was performed using a Leica DMi8 microscope equipped with a 25x/0.95 water immersion lens, Lumencor SpectraX LED lightsource, and a Hamamatsu Orca Flash 4 V3 sCMOS camera. The GCaMP6s calcium sensor was excited with 30% of the cyan LED of the SpectraX with a 488 nm excitation filter (≈ 12mW) and the calcium insensitive signal was excited with the 50% of the green/yellow LED through a 575/25 nm excitation filter (≈ 33mW) using a triple bandpass dichroic mirror in the filter turret (FF409/493/596-Di02-25×36, Semrock). The incident power of the excitation light was measured with a Thorlabs microscope slide power meter head (S170C) attached to PM101A power meter console. Emission was split with a Hamamatsu Gemini W-View with a 538 nm edge dichroic (Semrock, FF528-FDi1-25-36) and collected through two single band emission filters, 512/25 nm for GCaMP (Semrock, FF01-512/25-25) and 670/30 nm for mKate (Semrock, FF01-670/30-25), respectively. Both emission signals were split onto top/bottom of the image sensor, enabling differential exposure times optimized for imaging. Individual frames were acquired at 10Hz for 40-60 seconds (depending on worm movement) with an 88 ms and 50 ms exposure time, using the master pulse from the camera to trigger the light source through Hamamatsu HCImage software. Because it was impossible to resolve Ca^2+^ signals throughout the long DVA axon in moving animals, we restricted our curvature analysis to the posterior region close to the cell body.

###### Image analysis

Images were processed using custom built MATLAB routines to extract the mean intensity of the cell body as a function of body centerline curvature near the tail. Due to the omnipresent autofluorescent signal in the GFP channel, adaptive thresholding was able to separate the animal backbone from the background signal. First, the raw images from the GCaMP channel were binarized and eroded to find the skeleton of the worm in each frame. An iterative approach was chosen to prune the branches to get the longest path describing the centerline of the worm [73]. Next, a segment of the centerline enough to capture one bend of the worm in the tail region was chosen, which was further divided into two equal segments using three points. These three points were used to construct a triangle, and subsequently a circumcircle. Finally, the curvature is calculated at the middle point of the three points using the radius of the circumcircle by the formula *κ* = 1/R, where R is the radius of the circumcircle and *κ* is the curvature, and directionality of the curvature was determined by the sign of the tangential angle at the point where the curvature was calculated.

The neuron was labelled manually for the first frame in the mKate, calcium insensitive channel, and automatically tracked in subsequent frames, based on a local search in the vicinity of the location in the previous frames. The area, position and intensity was collected in the mKate channel and its position was mapped onto the GCaMP channel to extract the intensity in each frame. Finally, the calcium sensitive signal was divided by the insensitive signal after background correction (for Fig. S4) to obtain the ratio *R* which was normalized to the baseline ratio *R*_0_. The trough of the periodic signals was taken as *R*_0_. The background subtracted calcium traces were divided by the background subtracted mKate signal to yield the ratiometric intensity signal:

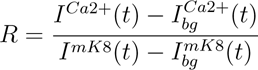

The fact that not all animals could be recorded with the same curvature change velocity and not all worms bent their bodies to the same extent, precluded the calculation of an average calcium signal *vs.* time. We thus transformed the periodic curvature *c* into a phase angle coordinate *θ* of 360 degrees [46] along the ventral-dorsal-ventral bending trajectory according to

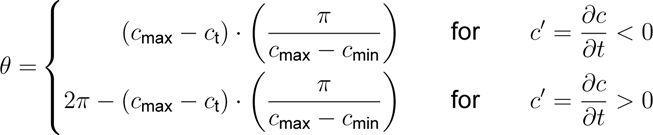

in which *c*(*t*) is the curvature at a given point in time, *c_max_* and *c_min_* is the maximum curvature and minimum curvature respectively, while *c^/^* is the first derivative of *c* with respect to time *t*. We arbitrarily assigned ventral to 0 and 360 degrees (start and end of the cycle) while dorsal corresponded to 180 degrees. The average calcium intensity ratio *R*/*R*_0_ was calculated and plotted against the phase angle.

##### Molecular biology and transgenesis

###### Construction of CRE driver lines

To drive CRE expression in target neurons, we constructed plasmids carrying CRE recombinase (synthesized from TWIST BioScience) and *F49H12.4p* for PVD [76], *nlp-12p* for DVA [21], *acr-5p* for B-type motorneurons [7] and *flp-22p* for SMD neurons [6]. All promotors were amplified using primers as listed below from genomic DNA isolated from N2 wt lab strains and cloned into a vector carrying codon-optimized CRE (TWIST BioScience) and synthetic introns designed according to [80] terminated with a *tbb-2* 3’ UTR using Gibson assembly. All clones were verified by Sanger sequencing.

###### Generation of *unc-70*(flox) using CRISPR

CRISPR/Cas9 genome editing was performed using previously described method with some modifications [81]. In brief, Cas9-crRNA-tracrRNA RNP complex together with repair templates (HDR) was assembled in IDT Nuclease-Free Duplex Buffer (30 mM HEPES, pH 7.5; 100 mM potassium acetate) and Mili-Q water. As in the co-CRISPR method we co-injected Cas9 complexes and repair templates targeting the marker gene *dpy-10* to incorporate the semi-dominant *cn64* allele. 20-30 young adult hermaphrodites were injected with the injection mix containing Cas9-crRNA-tracrRNA-HDR, and recovered onto individual plates. In some cases more than one crRNA were used. After 3 days post-injection, successful edits were identified based on the dominant *dpy-10(cn64)* roller phenotype and picked to individual plates to produce self-progeny. Mothers were then lysed and screened by PCR for the corresponding ed its using following primers as indicated in the following table. To generate the final genotypes, the CRE driver strains were bred with EG7944 [82] to mark the position of *unc-70* on chromosome 5 before crossing into *unc-70* floxed strain to establish the cell-specific CRE in the *unc-70*(floxed) background. Successful recombination of the *unc-70* locus was confirmed by PCR.

###### Conditional allele

We tagged twk-16 locus directly after the first ATG with worm optimized mScarlet, wScarlet, and a degron tag borrowed from the AID system, using Sunybiotech’s CRISPR services. We could identify wScarlet::TWK-16 expression in a single neuron in the tail (DVA) and two neurons in the head (probably AVK and ADE). Importantly, the fusion of TWK-16 with the AID::wScarlet tag did not cause any locomotion defects as compared to wt worms (Fig. S8E-G), indicating full functionality of the transgene.

###### Constitutive mutation

We removed 1672-bp from the genomic locus of *twk-16*, the region spanning the first two exons and the intron in between them. Two crRNAs were used to cut right before the start codon and in the intron region after the second exon, with a donor consisting of an ssODN containing the two 35-bp homology arms flanking the PAM sequence of the two crRNAs.

###### *mec-4p*::TRP-4

The *mec-4* promotor was amplified from pMH686 (gift of M. Harterink [83]) and cloned together using Gibson assembly with synthetic cDNA fragments produced by TWIST Bio-science, containing full-length TRP-4 (residues 1-1924) or truncated TRP-4 lacking the N-terminal domain with the ankyrin repeats (residues 1433-1924) to yield pNM2. The resulting plasmids were injected at 20 ng/*µ*l into GN692 [84] to yield MSB382 and MSB279 respectively.

###### DVA-Gal4 driver

The DVA-Gal4 driver was constructed using Gibson assembly of the backbone amplified from pHW393 (*rab-3p::GAL4-SK(DBD)::VP64::let-858* 3’UTR, addgene, plasmid # 85583) [28] and insert fragment containing *nlp-12p* amplified from pRD1 (*nlp-12p::CRE::tbb-2* 3’UTR) plasmid to yield pRD10. The resulting plasmid was injected at 5 ng/*µ*l together with *unc-122p::mCherry* as coinjection marker and integrated by UV irradiation using standard procedures.

###### DVA and TRN-TIR driver

The TRN-TIR driver was constructed using Gibson assembly of the backbone amplified from pUN1020 (*fln-1p::TIR::F2A::mCherry::H2B*, a gift from the Cram lab [27]) and insert fragment containing *mec-4p* amplified from pNM2 (*mec-4p::TRP-4::tbb-2* 3’UTR) plasmid to yield pNS30. The resulting plasmid was injected at 30 ng/*µ*l together with *unc-122p:mCherry* as coinjection marker. The DVA-TIR driver was constructed using Gibson assembly of the backbone amplified from pNS30 and insert fragment containing *nlp-12p* amplified from pRD1 plasmid to yield pNS43. The resulting plasmid was injected at 10 ng/*µ*l together with *unc-122p:mCherry* as coinjection marker.

###### Generation of the E2008K allele in UNC-70(TSMod)

pRD7 was constructed using Gibson assembly of the backbone amplified from pMK35(*pExprunc-70R8TSmodR9::unc-54* 3’UTR) and insert synthetic fragment (TWIST bioscience) containing E2008K mutation. The resulting plasmid was injected at 20 ng/*µ*l.

#### 1.5 Extrachromosomal array and integration

Extrachromosal arrays were generated by injecting the aforementioned amount of DNA into the appropriate strain and selecting for the F1 progenies with the co-injection marker. Three independent lines were generated whenever possible. The integration of the extrachromosomal array was performed using UV/TMP method. In brief, late L4-young adult animals carrying the array were picked onto a NGM plate without OP50. These animals were fed TMP at a final concentration of 50 µg/ml for 20 minutes. Then, they were UV irradiated for 30 seconds at 450 *mJ cm^−^*^2^ and expanded for 3-4 weeks before selection. Three independent integrated lines were recovered whenever possible.

#### 1.6 Primers and gRNA sequences

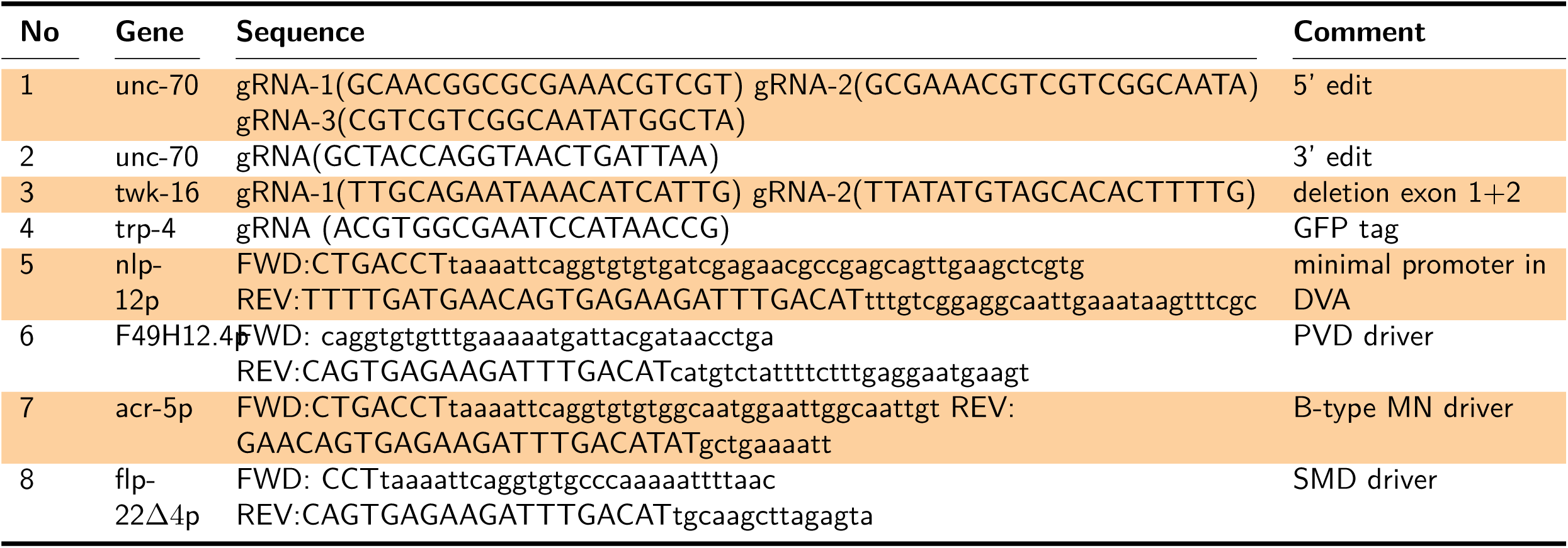

##### Primary culture and mechanical stimulation

###### DVA primary cell culture

Embryonic cell isolation was performed using a previously described method with some modifications [35]. Briefly, synchronized worms seeded onto peptone enriched plates and incubated at RT until the plates were populated with eggs. Then, the plates were washed off and the eggs were collected using milli-Q H2O. The worm and egg pellets were resuspended in the freshly prepared bleaching solution and rocked gently for 4-7 minutes, once 70-80% of the worms were lysed the reaction was stopped using egg buffer (118 mM NaCl, 48 mM KCl, 2 mM CaCl_2_, 2 mM MgCl_2_, 25 mM HEPES, pH 7.3 and an osmolarity of 340 mOsm.). The collected eggs were washed 3 times with fresh egg buffer. Then the eggs were separated using 30% final concentration of sucrose by centrifuging at 1200 rpm for 20 minutes. The separated top layers of eggs were collected in a new tube and washed 2-3 times with egg buffer. Then, the eggs were treated with chitinase (0.5 U/ml)) for 40 minutes to dissociate embryonic cells. The chitinase reaction was stopped using L15 medium. After chitinase treatment the embryos with the digested egg cells were passed through 25 G needle 10-15 times for dissociating into single cells. The dissociated cell suspension was filtered through 5 µm Durapore filter (Millipore). Then the single cell solution was centrifuged for 3 minutes at 3200 rpm and the pellet was resuspended in L15 medium.

Optical trapping chambers were based on two parallel glass surfaces spaced with a 50-*µ*m-thick polydimethylsiloxane (PDMS) layer (1:10 curing agent, Sylgard). A glass bottom Petri dish (BD, Wilco Glass, #1.5) was spin-coated for 1 min at 750 rpm [87] and cured for 1h at 65°C. A 1×1 cm square cavity was gently peeled off with a scalpel and the bottom dishes were then sterilized by UV irradiation for 1h. The cavity was then incubated with a 100 *µ*L drop of peanut lectin (Medicago, Sweden) diluted 1:10 into phosphate-buffered saline (PBS) for 20 min. After that, the plate was rinsed with L15 medium and a 150-*µ*L drop containing the cells was seeded. Cell density was 1.5 10^5^L*^−^*^1^ to ensure enough number of DVA neurons per plate, while keeping and appropriate space for the tether extrusion experiments. After 60 min, 0.5 mL of L15 medium was added and the cells were cultured at 25°C overnight. Fresh medium was changed (0.5 – 1 mL) every 24h. All experiments were carried out between one and four days post isolation.

###### Stimulation of DVA neurons cultured on elastic substrates

For the axonal deformation experiments, the neurons were directly seeded on the PDMS surface. The latter was obtained by spin-coating a bottom glass (Wilco Glass, #1.5) for 1 min at 750 rpm. The coated dish was then plasma-treated for 5 min (Plasma Surface Technology, Diener Electronic), UV-irradiated for 1h and incubated with 500 uL of peanut lectin (1:10 in PBS, Sigma Aldrich) for 24h. Axonal deformation was undertaken by indentation of the substrate with a 1-mm-thick microinjection needle held with a 4D-control micromanipulator (uMP-4, Sensapex). The needle tip was brought ≈5 mm away from the axonal tip and pushed vertically to induce substrate indentation. Substrate deformation was determined from the average perpendicular displacement at the middle of the axon. GCaMP activity was measured from the cell body and was normalized to the mKate, Ca^2+^-independent fluorescence signal (see below). A total of 25 cells were tested, 18 of which showed reproducible responses to substrate indentation.

###### Optical measurement of membrane mechanics and calcium activity

The optical tweezer (OT) platform (SensoCell, Impetux Optics, Spain) consists of a continuous wave laser (*λ*=1064 nm, 5 W nominal output power, Azur Light) steered with a pair of acousto-optic deflectors (AOD 1 and 2) and a force detection module that captures the forward-scattered light from the optical traps (Fig. S7A). This is mounted around an inverted research microscope (Nikon Eclipse Ti2) equipped with a spinning disk confocal microscope (Andor DragonFly 502, Oxford Instruments) on top of an active isolation table (Newport). The laser is directed onto a microscope objective (MO, 60x/NA=1.2, water immersion, Nikon) after being expanded with a telescope (lenses L1 and L2) to fill the MO entrance pupil, through the epifluorescence port. A short-pass dichroic mirror (D1) reflects the IR trapping beam and transmits both the excitation and emission light for fluorescence microscopy, as well as bright-field illumination (BF). To prevent from IR light leaking towards the detector, a neutral, shortpass filter (IR-F) was placed at the imaging optical path. The optical traps were positioned using LightAce software (Impetux Optics, Spain), in synchronization with a CMOS camera (BlackFly S, FLIR Systems) recording bright-field images of the sample. The force detection module of our optical tweezers platform operates by detecting light-momentum changes, after capturing the scattered trapping beam through an NA = 1.4, oil immersion collecting lens (CL), with a position-sensitive detector (PSD) placed at plane optically equivalent to the back focal plane (BFP) [88]. This allowed us to measure forces with no previous trap calibration and beyond the linear trapping regime, thus covering the full spectrum until the escape force of ≈200 pN, hence allowing working with lower laser power (250 mW), as compared to standard back focal plane interferometry [89]. The module enables the BF illumination (BF) to pass through, which is properly filtered to avoid leakage into the PSD.

###### Membrane tether extrusion

Prior to force measurements, red-fluorescent, 1-*µ*m polystyrene (PS) microspheres (FluoSpheres F8816, Thermo Fischer) were washed with cell culture medium by centrifuging at 10^4^ rpm for 3 min and added to the sample chambers at a concentration of 2 10^5^*µL^−^*^1^ and was then covered with a 1×1 inch coverglass (CG, #1.5, Ted Pella). The microspheres were captured in a 250-mW optical trap (laser power at the sample plane) and brought close to a DVA neuronal axon for 1 second before retraction to produce a stable membrane tether. The tethering force, *F_tether_*, was balanced with the trapping force, *F_OT_*, resulting in a change in the trapping beam momentum detected with the force detection module. Pulling routines were pre-defined with LightAce software (Impetux Optics) as follows (Fig. 5G): first, bead was contacted the axon and a pulled a distance of 10 *µ*m, reaching a peak in the tether tension value; second, the bead stopped for 10 s, letting the tether tension to relax down to a static value arising from the stored stress. Third, the bead was brought back to the initial position, releasing *F_tether_* to almost zero; finally, the bead stopped again for 10 s, letting the tether tension load up to a similar value. This routine was repeated three times for every DVA neuron tested at increasing velocities, 5, 20, 80 *µms^−^*^1^. Finally, every force measurement was compensated for initial momentum variation by subtracting a baseline trajectory taken on the same positions without a bead. The appearance of the Ca^2+^ transients is likely caused by the mechanical perturbation as opposed to local heating, as tether-free trials were unable to elicit reproducible changes (Fig. S6C,D). During each tether extrusion sequence, GCaMP and mKate fluorophores were simultaneously excited using the 488 nm and 561 nm laser lines of an Andor DragonFly Spinning Disk Confocal microscope, respectively. The excitation beam is corrected for non-uniformity and throughput using a Borealis Illuminator (BI, Andor, Oxford Instruments), before it passes through a quadband dichroic mirror (D2, 405-488-561-637 nm, Andor). After illuminating the sample plane, fluorescence emission is reflected at D2 and split with a 565-nm longpass dichroic mirror (D3) to image GCaMP and mKate emissions using F1: *λ*= 521 nm, F2: *λ*= 647 nm at two identical, back-illuminated scientific CMOS cameras (sCMOS 1&2, Andor, Oxford Instruments).

###### Data analysis of tether pulling

Peak (F*_peak_*) and storing (F*_base_*) force values were obtained from the trapping force signals during the tether extrusion experiments (Fig. S7B). Tension was calculated from the force according to Ref. [36]. GCaMP emission was measured both from the cell body (CB) and tether neck (TN) by setting a region of interest (ROI) with Fiji [91]. The background contribution was measured far from the neuron and subtracted from the intensity profile. The photobleaching trend was corrected by normalization over an exponential fit, providing the background subtracted and baseline normalized GCaMP signal (*F* /*F*_0_). Tension modulated calcium activity was observed in ≈30% of all extrusion events and is thus within the single molecule approximation [92]. To determine if the changes in neuronal membrane tension induced a significant increment in the GCaMP intensity, this was measured for a 3-second timeframe before and after the tether was extruded. Because the two, synchronized cameras recorded videos with a 10-frame rate, GCaMP intensity values were averaged over N=30 data points. The two intensity values (before and after pulling) were t-tested and thresholded within a *p*-value of *<*0.01. When significant, the pulling events were classified into GCaMP-increasing (Δ*I >* 5%) and GCaMP-decreasing (Δ*I <* 5%). To rule out that the Ca^2+^ transient were caused by heating of the trapping laser, we carried out a series of measurements on wt DVA neurons in the absence of a membrane nanotube tethering the trapped microbead and the neuronal axon (without prior contact of the bead).

#### 1.7 Monte Carlo simulation of force-gated ion channel ensembles

To capture the dynamics and the statistical behavior resulting from the stochastic activation of an ensemble of mechanosensitive ion channels, subjected to a mechanical force, we set up a continuous time Markov chain Monte Carlo simulation [93]. We choose to model a pair of mechanosensitive ion channels, which we conceptualize as an excitator, sodium or calcium conductive in channel and an inhibitory, potassium or chloride conductive ion channel. Our model is agnostic of the force transmission pathway and does not differentiate between membrane and cytoskeletal force delivery. To simulate the behavior in absence of external noise, we assumed that each channel acts independent, activities are uncoupled, and each channel is characterized by an open and a closed state that is separated by a potential barrier with height *E* (Fig. S7). The lifetime of each state dependents on the height of the energy barrier separating the closed from the open states and the loading conditions. Opening is driven by thermal fluctuations, and, as a result, is a stochastic process. Application of force to the channel tilts the energetic landscape, thus reducing the energy barrier that separates the closed from the open state by an amount *F* · γ, in which *γ* is the distance to the transition state [38]. If a load is applied to the channel for durations that are much shorter than the intrinsic lifetime of the closed state, the channel resists opening. Importantly, channels do not confer resistance to force on timescales that are larger than the intrinsic lifetime of the particular closed state [95, 96]. In agreement with previous data on whole cell recordings from TRP-4 [39], we assumed that the excitatory channel activates at the onset and the offset of the force. Such behavior is consistent with a strain-rate sensitivity [98], thus, we model the channel sensitive to the first derivative of the force, 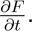. The forward transition rate was model using the modified Evans-Bell model of time-dependent bond-strength, as determined by the force-rate or loading rate *r_f_* [96]. Loading rate was calculated from the stiffness of the ankyrin domain [99] multiplied by the pulling velocity in the experiment. We start the simulation with all states closed, and are interested in the evolution of the ensemble to the open state.

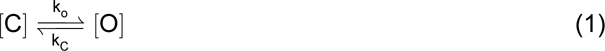

The lifetime of the closed state is governed by the spontaneous opening constant 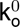 according to

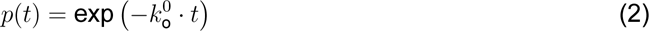

For an open channel, the probability of finding the channel open after time *t* decays exponentially and will spontaneously revert back to the closed state stochastically if the random sampling parameter 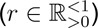 is smaller than *p*(*t*). Thus, if a channel is in the open state at time *t*, the probability of finding it in the open state *t* + 1 decreases *e*-fold:

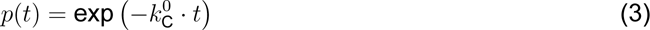

Force sensitivity is achieved by applying Bell’s model to the forward rate constant. We likewise assume that the channel cannot sustain the open state as long as force is acting. This assumption has the physical manifestation in a force-transmission pathway through a weak protein-ligand interaction (slip bond). After time *t*, we apply a force to the channels. Thus, the probability of a closed channel responding to the external forces changes to

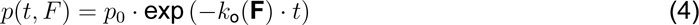

in which

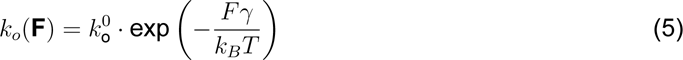

Evan’s modification for a finite loading rates was implemented to capture the strain-rate dependence of the Calcium channel (TRP-4), known to respond to the change in force.

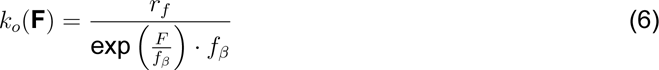

with 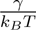 as the force scale.

We implemented the simulation in R, with a timestep of 1e-5 s and the following kinetic constant. Inhibitory Channel: 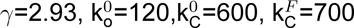

Excitatory channel: 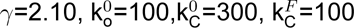

The physical representation of the values 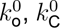 correspond to the spontaneous opening constants. For 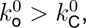 ion channel remain statistically open, otherwise they spent more time in the closed state on average. Without a lack of generality, the concept can be applied for lipid bilayer tension-gated ion channels in which the free energy profile of the energy landscape is altered by a external tension *σ* according to ΔΔ*G* = −Δ*G* − σ · Δ*A*, in which Δ*A* equals to the increase in cross sectional area of the gated ion channel, e.g. Δ*A*=4.7nm^2^ for TREK2 [17]. Thus, the tension dependent *k_o_* conforms to 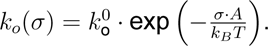 It can be readily seen that without an increase in cross-sectional area, the open state is not preferred. Finally, the average current was calculated by *I* = *cNP_o_*, where *c* is the single-channel current taken from the literature (TRP-4, 18pA; 140pS [101]; K2P, 13pA; 90pS [102]), *N* the number of channels, and *P_o_* the average probability of finding the channel open derived from the simulations. Under assumption of a high input resistance typical for *C. elegans* neurons [60], the K signal was then subtracted from the Ca signal to yield a macroscopic ‘observable’, distantly related to a Calcium signal. The picture that is emerging from this simulation is that ventral DVA activity emerges in part from TRP-4 activation under compression and the suppression of ‘stretch’ currents under dorsal side. Whereas this describes a plausible explanation for our findings, two other possible scenarios could give rise to the observed ventral activity *in vivo*: 1) TWK-16 and TRP-4 both activate under tension, and close with different rates such that a remaining Ca^2+^ activity is visible during ventral bouts (Fig. S7F); and 2) TRP-4 is constitutively active and only modulated by TWK-16 leading to Ca^2+^ suppression during tension (Fig. S6G). The combined results from our *in-vitro* (Fig. 5), *in vivo* (Fig. 6) and *in-silico* (Fig. S6e-g) experiments favor a scenario in which mechanosensitive TWK-16 activity suppresses stretch-induced depolarization. Because TRP-4 is a pore-forming sub-unit of a mechanosensitive ion channel that activates at the force onset and offset [39] we should expect DVA activity during dorsal AND ventral bends, but we exclusively recorded Ca^2+^ increases during force relaxation/offset *in vitro* and compressed axons during ventral bends *in vivo*. In absence of TWK-16, however, the biphasic TRP-4 activity is unveiled.

#### 1.8 Neuromechanical model

Without exception, all parameters and assumptions have been reproduced as outlined in Ref. [44]. In short, the framework consists of a 2D structural skeleton composed of 46 segments and 98 discrete joints distributed on the ventral and dorsal sides. These joints are vertically connected by incompressible rods and lateral connections embody passive forces modeled as a Kelvin-Voigt material with a spring and a dashpot in parallel owing a constant material property. Diagonal elements, connecting neighboring joints on opposing sides, represent the effect of pressure and ensure volume conservation. In parallel to the passive force, ‘muscle’ forces are computed as an active Kelvin-Voigt material with varying spring constants and viscosities, whose parameters are slaved to neuronal state variable of the motor circuit (active, inactive). For details see ref [44].

The minimal motorcircuit was embedded into that active/passive framework, consisting of a pair of excitatory (DB,VB) and inhibitory motorneurons (DD,VD). The current within the motor ventral motorcircuit has contribution from stretch receptors, in addition to the input current from AVB and motor current driving muscle actuators. The current in dorsal motor neuron does not contain adjustments from the stretch receptor. The modified the model according to our experimental data: We eliminated dorsal stretch receptor currents 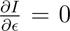 (with ∊ = *dc* as the strain) and inverted the relationship of the stretch receptor current on the ventral side and deliver it to B-class motor neurons on the dorsal side. No other stretch receptors were considered on the ventral side. In this model a compression on the ventral side generates a polarising response, which results in polarising current flowing into the B-class motor neurons on the dorsal side, and *vice versa*, the stretch on the ventral side generates a hyperpolarizing response. The DVA neuron is implemented indirectly, as a compression/stretch sensor, sending either positive or negative current to DB neurons. (A similar effect can be achieved by modifying the firing rate of DVA, which is non-zero at resting length). The factor modifying the effective stretch receptor activation function (*−***S**) was implemented in the original [44, Eq. 11]:

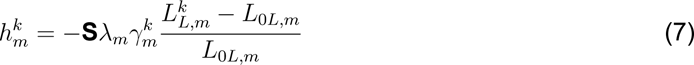

where **S** scales the effective stretch receptor activation function. This may be understood as decreasing the number of effective stretch/compression sensitive ion channels, or a failure to recruit active ion channels (equivalent to a failure in force transmission).

#### 1.9 Statistics

Statistical analyses were performed in R and Igor. To construct and visualize the 3D distributions of the modes (*a*_1_, *a*_2_, *a*_3_), the 3D kernel density estimate of the first three modes was calculated in R using the ks package [105] with an unconstrained plug-in selector bandwidth. We choose to indicate the 10, 25 and 50% contours of the highest density regions in the manifold and the 2D projections of the floating data cloud along the corresponding planes. To compare two different data sets and test for the null hypothesis that the two kernel density functions are similar, we resampled the highly oversampled population by bootstrapping to avoid spurious significance due to long tailed outliers. The resampling and testing was performed 1000 times to yield a distribution of *p*-values which is displayed as a violin plot summarizing each figure of locomotion data. Importantly, the resampling does not lead to significant discrimination of the downsampled and original dataset within the same population. Alternatively, the local density between two distributions was tested using a binned kernel density estimator. For this, we used the `kde.local.test` function from the ks package in **R** [105] to a) converted all n data points of each distribution to counts on a 100×100×100 binning grid and embedded into a matrix C, b) evaluate the kernel function at these grid points to embed them into a matrix *K* and c) then obtain the binned density estimator *f* from a sequence of discrete convolutions of *C* and *K*. The exact formulation of that procedure and a detailed presentation of the algorithm can be found in ref. [106]. For all grid points in which *p >*0.01, we accept the null hypothesis that the two densities at this grid point for the two distributions is the same. For all other grid points, a polarity is assigned and plotted as two different colors in a 3D voxelgram (e.g. Fig. 1D), indicating that *x*1 *> x*2 or *x*1 *< x*2. Swarm plots and estimation statistics have been calculated using the methods described in ref. [107].

## 2 Supplementary Videos

**Video S1: Locomotion behavior in wt and unc-70 mutants** Representative video of wildtype (N2) and *unc-70(e524)* (CB524) mutant animal. Acquired at 25Hz

**Video S2: Crawling behavior of conditional unc-70 alleles.** Representative video of conditional CRE/loxP mutant strains in the order as they appear in Figure 3. Scale bar = 300*µ*m, acquired at 25Hz. *unc-70*(floxed) is the control animal without CRE expression denoting the background for all other genotypes. Pan-neuronal, *rgef-4p*::CRE; BWM, body wall muscle restricted *myo-3p*::CRE; D-type MN, GABAergic motorneuron directed *unc-25p*:CRE; B-type MN, cholinergic forward motorneurons directed *acr-5p*::CRE; A-type MN, cholinergic backwards motorneurons directed *unc-4p*::CRE; TRN, touch receptor neuron specific mec-17p::CRE; SMD, SMD-directing *flp-22*Δ *4p*::CRE; PVD, PVD-directing *F49H12.4p*::CRE; DVA, DVA-specific *nlp-12p*::CRE in the *unc-70*(floxed) background.

**Video S3: DVA Calcium activity in unc-70 mutants** Representative video of DVA calcium activity in wildtype (left), *unc-70(e524)* (middle) and conditional DVA CRE/loxP mutant strains (right). Upper panel shows the calcium sensitive GCaMP6s, lower panel a calcium-insensitive mKate as a movement and defocussing control. Playback speed, 30 frames/s.

**Video S4: DVA responds to substrate deformation** False color labeling of a GCaMP6s expressing DVA neuron cultured on PDMS, subjected to a mechanical deformation. Yellow shows the calcium sensitive GCaMP6s, magenta a calcium-insensitive mKate as a movement and defocussing control. Scale bar = 5*µ*m. Acquired at 10Hz.

**Video S5: Calcium activity in DVA during dynamic membrane tether extrusion** Representative video of DVA neuron in the dynamic optical trapping assay. Scale bar = 5*µ*m. Acquired at 10Hz.

**Video S6: Locomotion behavior of conditional twk-16 mutant animals** Representative video of a TWK-16::AID animal in presence of 1mM auxin; left animal without (MSB555) and right with (MSB526) DVA::TIR expression. Scale bar = 300*µ*m. Acquired at 25Hz.

**Video S7: Compression induced proprioceptor current coordinates locomotion behavior** Left: Animation derived from the results of the neuromechanical model for input parameters giving rise to wildtype-like animal locomotion pattern implementing DVA as a compression sensitive proprioceptor. Right: Same model with lower sensitivity to curvature induced compression current in DVA, representing *trp-4* and *unc-70* mutations.

## 3 Supplementary Figures

**Supplementary Fig. S1.**
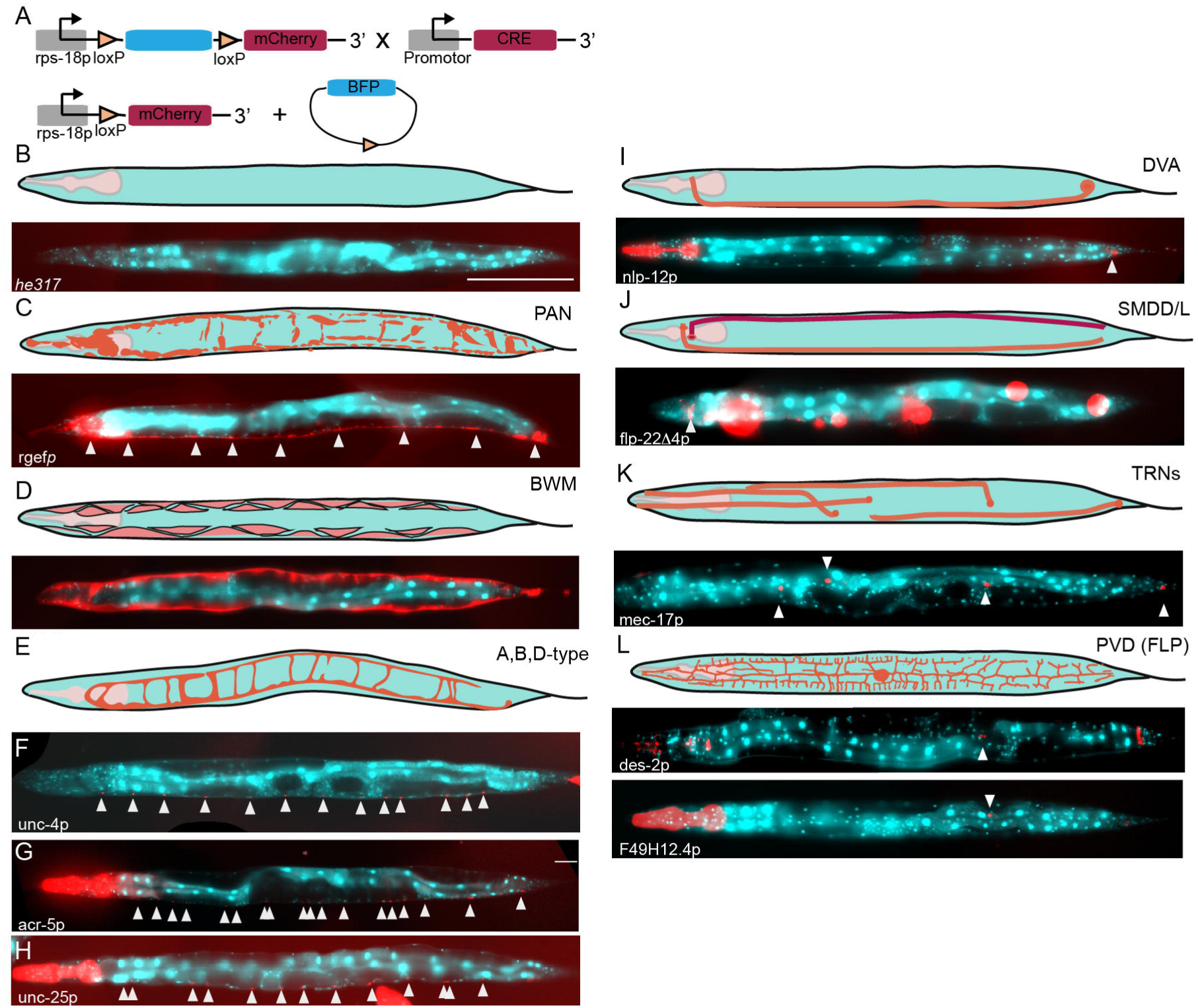
Reporting CRE recombination efficiency. **A** Strategy of the CRE recombination reporter. A floxed tagBFP with a nuclear localization signal (NLS) under the control of the ubiquitous *rps-18* promoter is visible before recombination. After CRE expression in specific cells and tissues, the tagBFP gets excised and brings an NLS::mCherry construct under the control of the *rps-18p*, enabling the identification of targeted cells. For details and number of animals investigated see Table S2. **B** Schematic of the predicted pattern and representative picture of a reporter animal without CRE expression showing only BFP expressing cells. **C** Schematic of the predicted pattern and representative picture of a panneuronal CRE activity under *rgef-1p*. **D** Schematic of the predicted pattern and representative picture of a CRE activity in body wall muscles under *myo-3p*. **E** Expected pattern for motorneurons. **F** Representative picture of a CRE activity in A-type motorneurons under *unc-4p*. The red dot in the tail is due to lin-44::DsRed coninjection marker. **G** Representative picture of a CRE activity in B-type motorneurons under *acr-5p*. The pharyngeal signal is due to myo-2p::Cherry coinjection reporter. **H** Representative picture of a CRE activity in D-type motorneurons under *unc-25p*. The pharyngeal signal is due to *myo-2p::Cherry* coinjection reporter. **I** Schematic of the predicted pattern and representative picture of a CRE activity in DVA neuron under *nlp-12p*. The pharyngeal signal is due to myo-2p::Cherry coinjection reporter. **J** Schematic of the predicted pattern and representative picture of a CRE activity in SMD under *flp-22*Δ*p*. The six red spots belong to *unc-122p::RFP* coinjection reporter. **K** Expected recombination pattern for touch receptor neurons (TRNs). Representative picture of a CRE activity in TRNs visible in four (on the left side) out of the six neurons. The right side is not imaged. **L** Schematic of the predicted pattern and representative picture of a CRE activity in PVD under the control of the *des-2p* and *F48H12.2p*.

**Supplementary Fig. S2.**
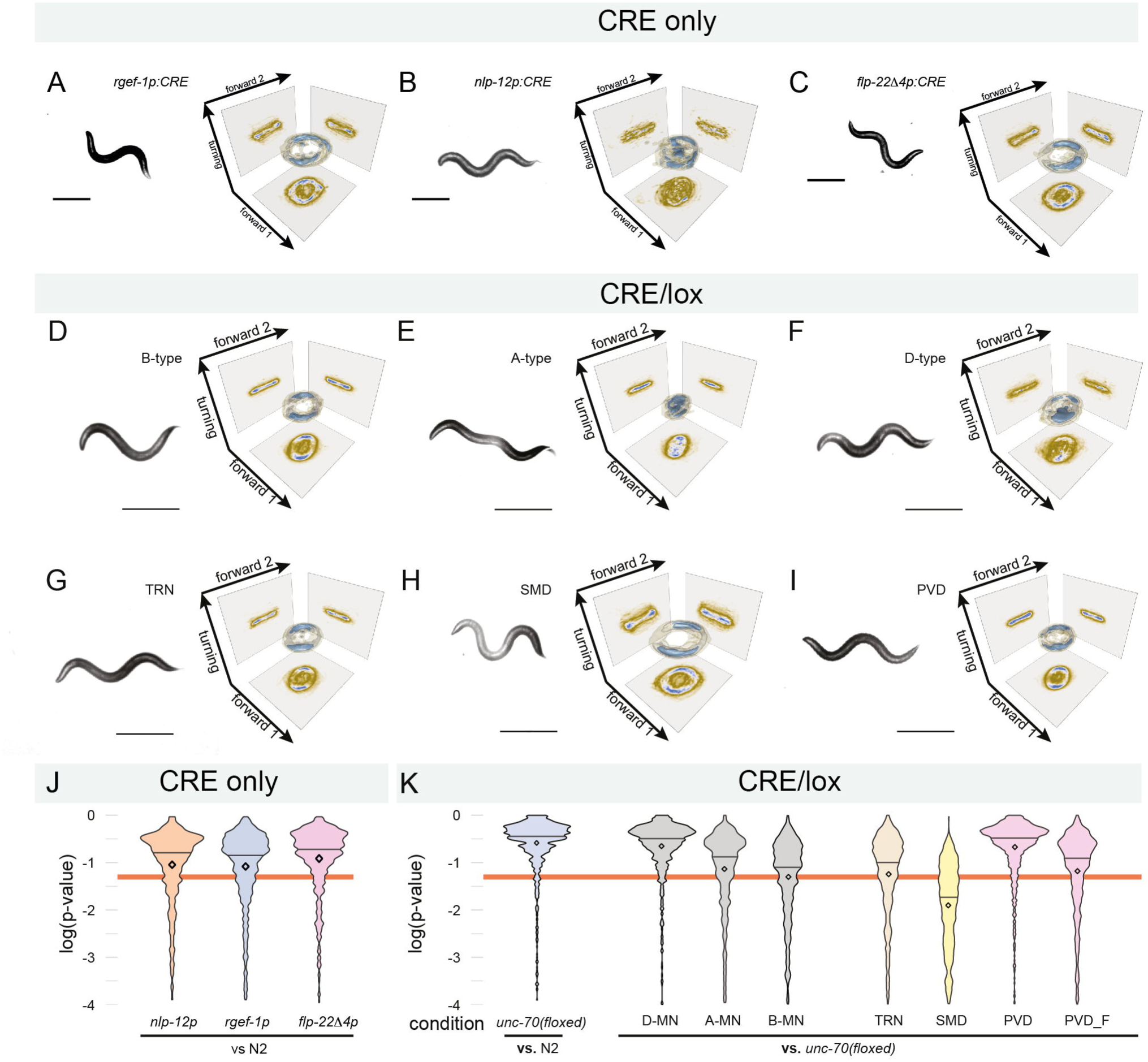
DVA-specific mutation of the spectrin network causes abberrant body postures. **A-C** Still image and the corresponding 3D eigenworm orbit for (**A**) pan-neuronal CRE, (**B**) *nlp-12p*::CRE and (**C**) *flp-22*Δ*4p*::CRE expressing animals. Only CRE drivers are shown that showed a phenotype in combination with the unc-70(flox) allele. D Distribution of *p*-values for the hypothesis test H0 that the two data sets indicated in the graph are sampled from the same population. Orange line indicates *α*=0.05 level of significance, black diamond represents the mean and horizontal line the median *p*-value of the distribution. F-K Representative still image and the corresponding manifold in the three dimensional eigenworm space for (**F**) D-type, (**G**) B-type, (**H**) A-type motorneurons, (**I**) in TRNs, (**J**) SMD, and (**K**) PVD. **L,M** Distribution of *p*-values for the combinations indicated in the figure. Orange line indicates *α*=0.05 level of significance.

**Supplementary Fig. S3.**
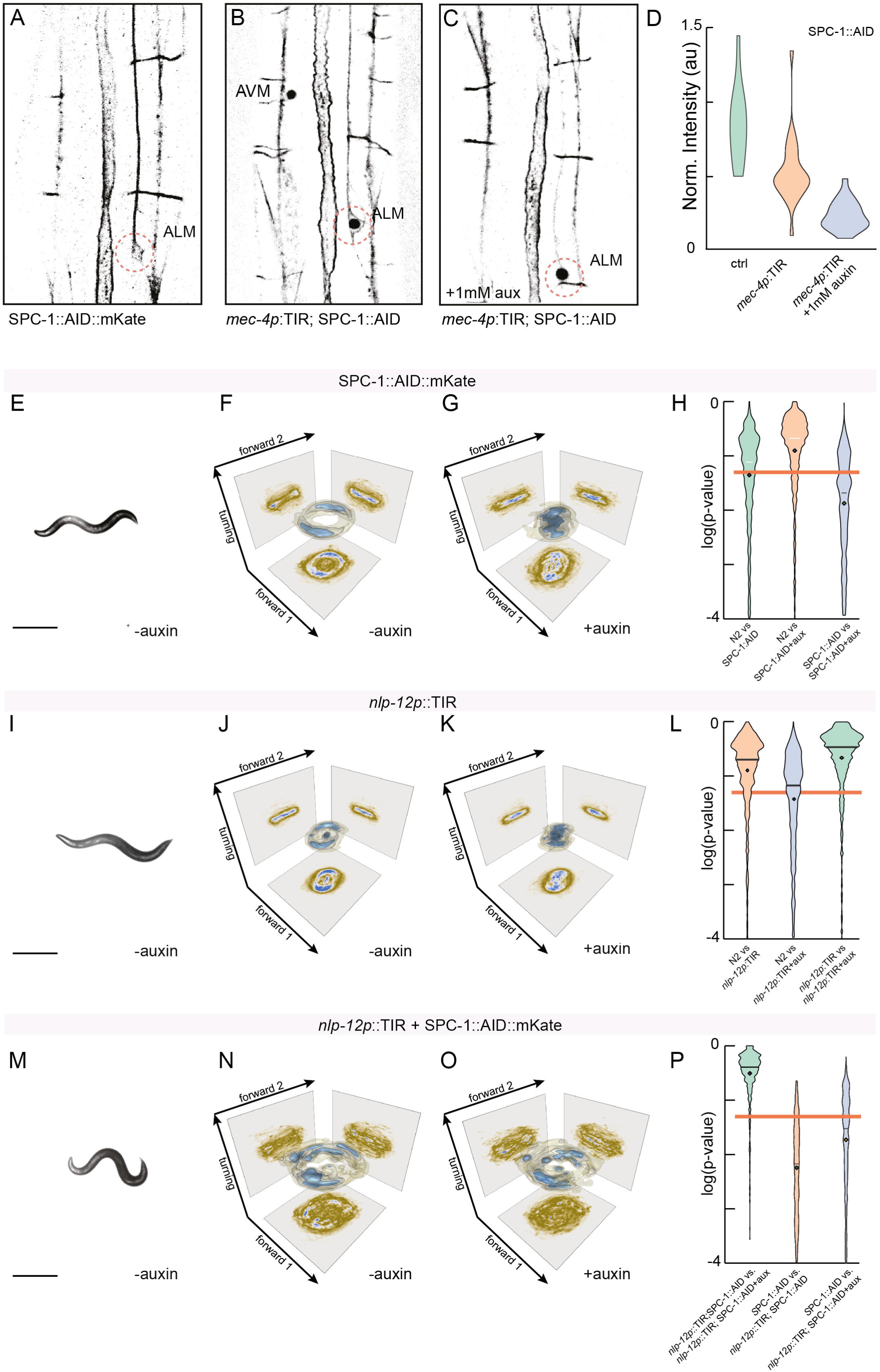
SPC-1 shares function during locomotion in DVA with UNC-70. **A-C** Representative images of an (**A**) UN1823 (SPC-1::AID::mKate) expressing control animal (without TIR ligase) and (**B**) MSB453 (*mec-4p*::TIR->SPC-1::AID::mKate) with nuclear mCherry localization indicating TIR expression in TRNs in absence and (**C**) presence of 1mM auxin. Due to the overlap of DVA axons with other neurites in the ventral nerve chord, we choose to estimate the effect in TRNs. **D** Quantification of the neurite intensity of TRNs without TIR (ctrl, N=17 animals) and with TIR ligase in absence (N=19) and presence of auxin (N=20) in the SPC-1::AID::mKate background, normalized by the intensity of motorneuron commissure (that do not express the TIR ubiquitin ligase). **E-G** Representative snapshot of a (**E**) SPC-1::AID::mKate animal without TIR ligase and the corresponding quantification of its behavior in (**F**) absence and (**G**) presence of auxin. **H** Distribution of *p*-values as described above for the combinations indicated in the figure. **I-K** Representative snapshot of a (**I**) MSB503 animal expressing TIR exclusively in DVA (*nlp-12p*::TIR::F2A::H2BmKate) and the corresponding quantification of its behavior in (**J**) absence and (**K**) presence of auxin. **L** Distribution of *p*-values for the combinations indicated in the figure **M-O** Representative snapshot of a (**M**) MSB464 animal expressing TIR exclusively in DVA together with the SPC-1::AID::mKate degron and the corresponding quantification of its behavior in (**N**) absence and (**O**) presence of auxin. **P** Distribution of *p*-values for the combinations indicated in the figure. Note, due to the auxin-independent TIR activity, the addition of 1 mM auxin does not further increase the auxin-independent loss of coordination.

**Supplementary Fig. S4.**
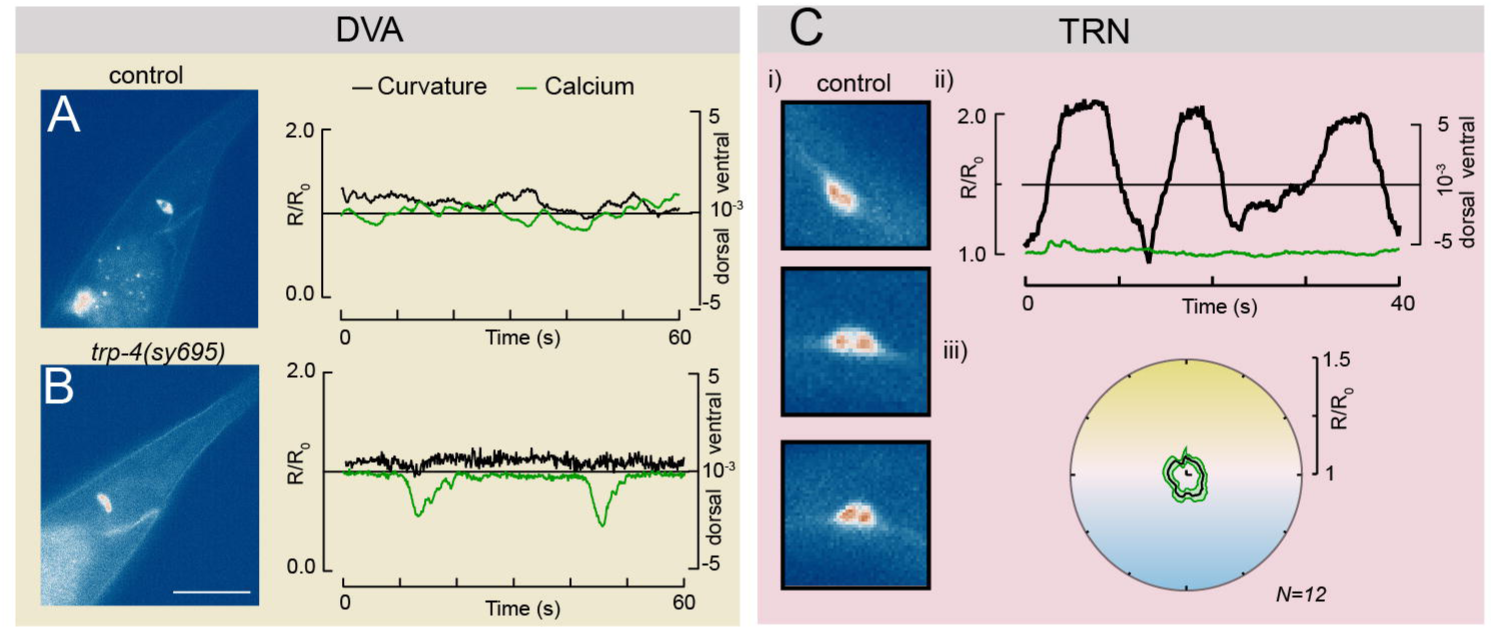
Cell autonomous calcium activity in immobilized animals and TRNs. **A,B** Single still images of a tail from (**A**) control and (**B**) *trp-4* mutant animals and the quantification of curvature and spontaneous calcium activity displayed as normalized GCaMP6s/mKate ratio on the left. Scale bar = 50*µ*m. Images and traces representative for 12 and 11 animals, respectively. **C** Calcium activity in control PLM touch receptor neuron (without ectopic TRP-4 expression). i) Representative images of the calcium-sensitive GCaMP6s expressing in PLM cell body under ventral, neutral and dorsal body bends. False colored Vik palette. ii) Curvature and ratiometric calcium signal plotted against experimental time, showing little to no modulation of calcium transients under modest curvatures.iii) Quantification of the average GCaMP6s/tagRFPt ratio as a function of the phase angle of the dorso-ventral body curvature.

**Supplementary Fig. S5.**
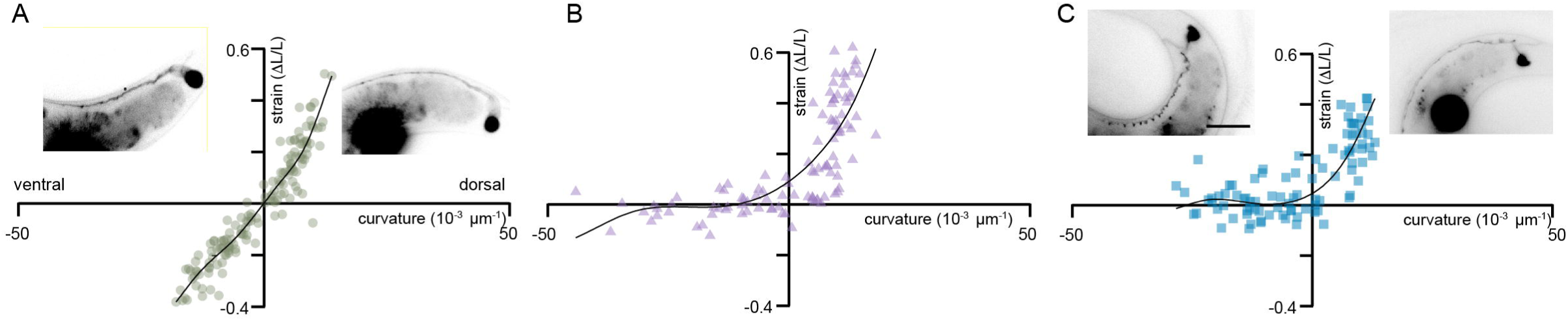
DVA is under compression during ventral bends. **A-C** Normalized length change in DVA vs body curvature in (**A**) wildtype, (**B**) *unc-70(e524)* and (**C**) DVA::unc-70(0) animals. Black line indicates the running average of the individual datapoints shown in colored circles with the slope corresponding to the compliance of the neuron. Representative morphologies corresponding to DVA under compressive and tensile body curvatures are depicted in the inset epifluorescence micrograph of a DVA::mKate expressing animal.

**Supplementary Fig. S6.**
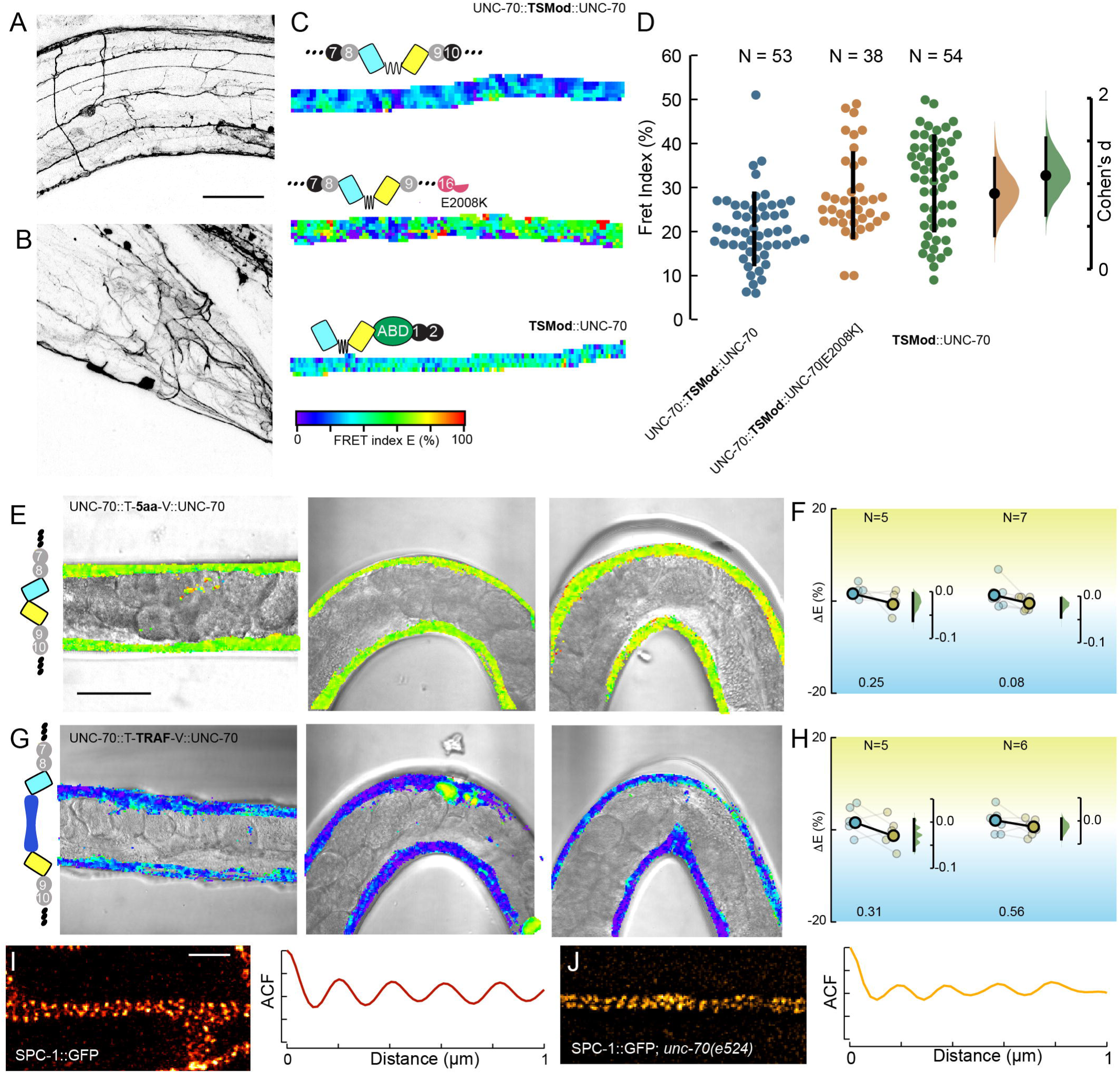
*β*-spectrin organization and mechanics. **A** Maximum intensity projection of high resolution confocal images of N-terminal *β*-spectrin fusion under the control of the endogenous 2kB *unc-70* promoter used for TSMod expression (for details about construction, see Ref. [19]), showing predominant expression in neurons and faint expression in muscles. Scalebar = 20 *µ*m. **B** Posterior image of the same animals as in (**A**). **C** Representative ROIs of different neurons (of untracked identity) expressing the tension sensor module embedded into wildtype and E2008K mutant *β*-spectrin compared to the N-terminal no force control. **D** Swarm plot of the average FRET efficiency per neuronal ROI analysed for the three transgenes. The Cummings plot on the right indicates the bootstrapped distribution of the Cohen’s d as calculated from the mean difference taken from 5000 trials divided by the combined standard deviation comparing control vs E2008K and control vs N-term. The vertical black bar indicates the 95% confidence interval. **E,F** FRET measurement in a transgenic line expressing a constitutive high FRET construct (mTFP-5aa-mVenus) embedded between repeats 8 and 9. **G,H** FRET measurement in a transgenic line expressing a constitutive low FRET construct (mTFP-TRAF-mVenus) embedded between repeats 8 and 9. **I,J** Representative STED images and autocorrelation of (**I**) SPC-1::GFP expressing neurons and *unc-70(e524)* mutant animals expressing SPC-1::GFP [29]. Scalebar=1*µ*m

**Supplementary Fig. S7.**
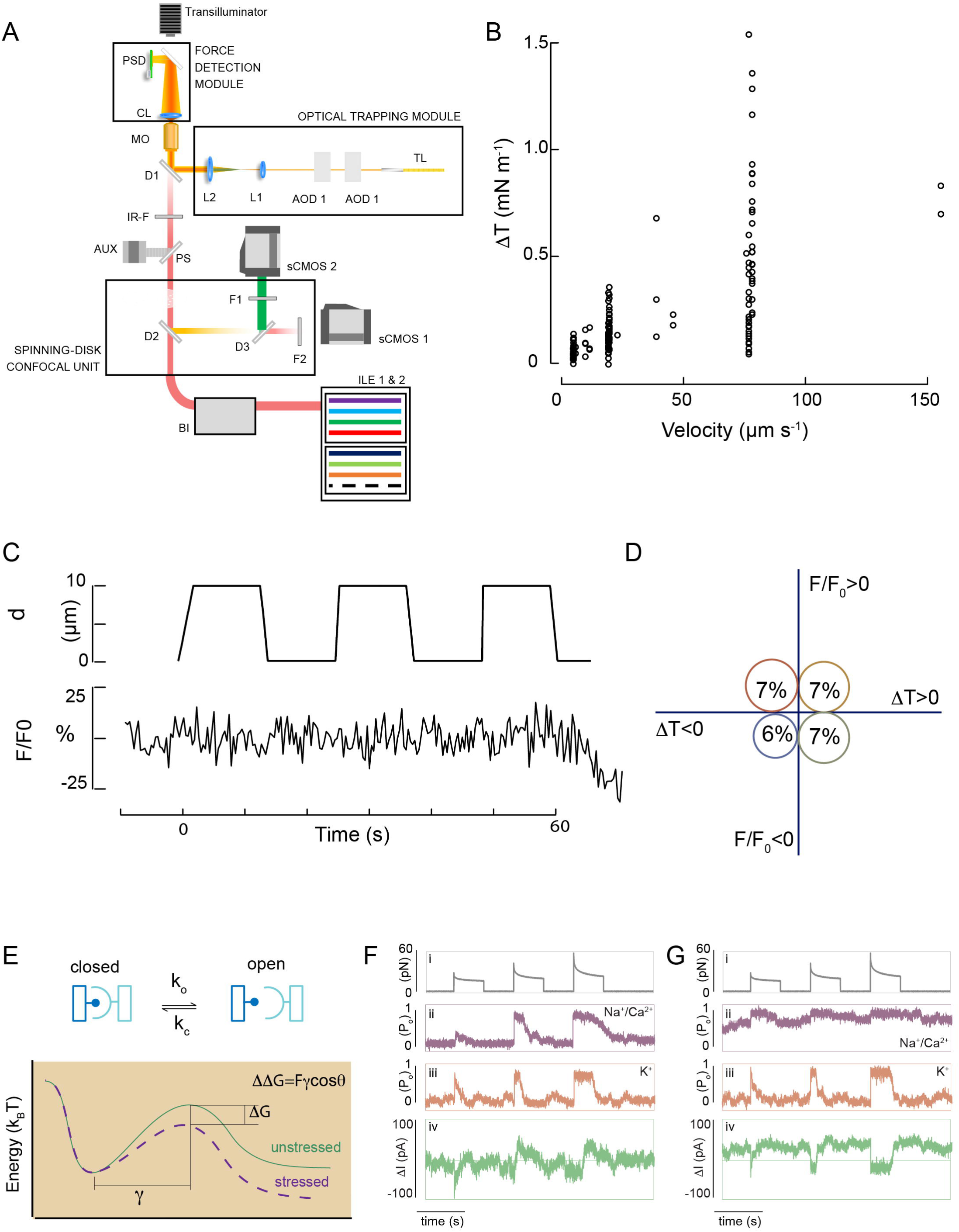
Dynamic tether force spectroscopy of isolated proprioceptor neurons. **A** Schematic of the set-up combining spinning-disk confocal microscopy and optical trapping. (ILE, integrated laser engine; BI, Borealis Illuminator; D, dichroic mirror; F, filter; IR-F, IR filter; CL, optical tweezers collecting lens; PSD, position-sensing detector; TL, transmitted light source; L, lens; AOD, acousto-optic deflector; LS, trapping laser source; AUX, eyepiece camera). **B** Membrane tension (Δ*T* = *T*_Peak_ − *T*_base_) gradient measured for each extrusion event as a function of velocity. Tension was derived from the different between the peak force and the plateau force of the tether extrusion experiments according to 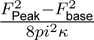 with *κ* = 2.7*e −* 19Nm as the bending rigidity of the axonal membrane [36]. **C** Representative displacement, force and bleach-corrected calcium trace for the tether-free no-force control. **D** GCaMP variation versus tension gradient bubble plot for the tether-free negative control. N=108 events on n=36 cells. **E** Schematic of how force tilts the hypothetical 1-D energy landscape, with the location of the transition state *γ* separating the closed and open conformation of the mechanosensitive ion channel. **F** Simulation of a cation (purple) and K+ selective (orange), mechananosensitive ion channel that solely respond to the force onset and close with different kinetics 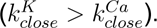 The green trace resembles the Calcium dynamics under the assumption of an unchanged input resistance and single channel conductance. **G** Simulation of a constitutively active cation selective ion channel (purple) and a mechanosensitive K+ ion channel (organge). The forced activity of the K+ channel modulates the observable (combined open probability, green trace).

**Supplementary Fig. S8.**
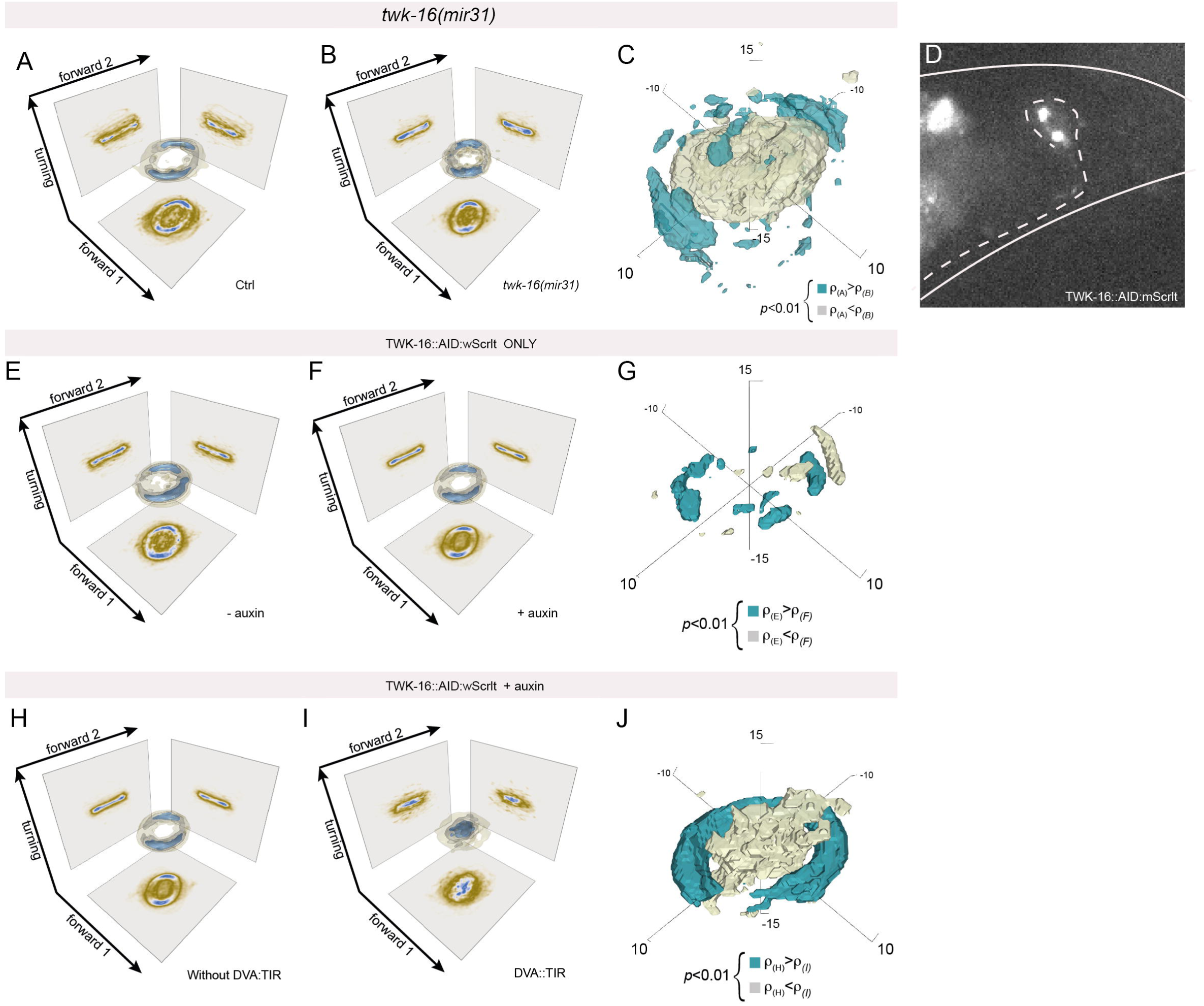
Suppression of DVA activity through TWK-16 modulated locomotion behavior. **A-C** 3D density estimate for joint probability distribution of the two forward and turning modes in the eigenworm space for (**A**) control and (**B**) *twk-16(mir31)* mutant animals and (**C**) the statistically significant differences in the local density functions *ρ_ctrl_* and *ρ_twk−_*_16_. Blue voxels indicate higher local density for ctrl, beige voxels indicate higher density for *twk-16* on the *α*=0.01 level. D Micrograph of a DVA neuron expressing TWK-16::AID::wScarlet, representative for *>*20 animals. E-G Corresponding 3D density functions for TWK-16::AID control animals in (**E**) absence and (**F**) presence of auxin, and the statistically significant differences in the local density functions on the *α*=0.01 level. Note, the discontinuity in the distribution of the p-values, indicates high similarity of the 3D probability functions. H,I 3D density functions for TWK-16::AID::wScarlet animals (**H**) WITHOUT TIR expression and (**I**) WITH DVA restricted TIR expression. Both distribution were recorded in presence of auxin. J Statistically significant differences between the local density functions *ρ_H_* and *ρ_I_* displayed in panel and (**I**).

## 4 Supplementary Tables

**Table S1.** Locomotion data. Strains used for the data acquisition of animal locomotion behavior

| Strain | genotype | condition | Fig. | Neuron | worms | frames |
| --- | --- | --- | --- | --- | --- | --- |
| N2 |  |  | 1 |  | 41 | 75985 |
| CB524 | unc-70(e524) |  | 1 |  | 20 | 20000 |
| MSB187 | unc-70 (mir6mir16) V; tmls1070 | CRElox | S2 | a-type MN | 21 | 35933 |
| MSB535 | mirls37; unc-70 (mir6mir16) V; | CRElox | S2 | b-type MN | 20 | 44077 |
| MSB239 | tmls1087; unc-70 (mir6mir16) V | CRElox | 2 | BWM | 19 | 38600 |
| MSB186 | unc-70 (mir6mir16) V;tmls1072 | CRElox | S2 | D-type MN | 19 | 25702 |
| MSB160 | tmls777; unc-70 (mir6mir16) V | CRElox | 2 | PAN | 20 | 20078 |
| MSB536 | mirls42; unc-70 (mir6mir16) | CRElox | S2 | PVD | 20 | 42268 |
| MSB450 | heSi208; hrtSi27 V;ll;<br>unc-70(mir6mir16)V | CRElox | S2 | PVD | 11 | 19941 |
| MSB295 | mirEx98; unc-70 (mir6mir16) V | CRElox | S2 | SMD | 19 | 31331 |
| MSB424 | heSi317; hrtSi99;<br>unc-70(mir6mir16) | CRElox | S2 | TRN | 10 | 21125 |
| MSB188 | mirEx13; unc-70(mir6mir16) | CRElox | 2 | DVA | 38 | 62307 |
| MSB539 | mirls43;unc-70(mir6mir16) | CRElox | 2 | DVA | 17 | 34406 |
| MSB225 | trp-4(sy695);unc-70(e524) |  | 4 |  | 20 | 6694 |
| MSB250 | trp-4(sy695);unc-70(DVA) |  | 4 |  | 6 | 11049 |
| TQ296 | trp-4(sy695) |  | 4 |  | 23 | 28577 |
| MSB115 | unc-70(mir6mir16) | no CRE | 2 |  | 10 | 35646 |
| GN717 | trp-4(ok1605) |  | ns |  | 21 | 39086 |
| MSB516 | mirls43; he317 | no floxP | ns | DVA | 17 | 37668 |
| MSB66 | mirEx13 | no floxP | S2 | DVA | 5 | 11147 |
| FX14125 | tmls777 | no floxP | S2 | PAN | 12 | 25008 |
| MSB340 | mirEx98 | no floxP | S2 | SMD | 17 | 23700 |
| MSB464 | mirEx194;<br>spc-1::degron::mKate2 | TIR+AID+aux | S3 |  | 15 | 17840 |
| MSB464 | mirEx194;<br>spc-1::degron::mKate2 | ctrl | S3 |  | 15 | 16872 |
| UN1823 | spc-1::degron::mKate2 | AID +<br>auxin | S3 |  | 15 | 20574 |
| UN1823 | spc-1::degron::mKate2 | AID ctrl | S3 |  | 15 | 20666 |
| MSB503 | mirEx197 | TIR ctls | S3 |  | 15 | 22151 |
| MSB503 | mirEx197 | TIR +<br>auxin | S3 |  | 15 | 21781 |
| MSB555 | twk-16(syb2541) | AID ctrl | S8 |  | 10 | 21323 |
| MSB555 | twk-16(syb2541) | AID +<br>auxin | S8 |  | 16 | 34971 |
| MSB521 | twk-16(mir31); mirls19; syls423 |  | 6, S8 |  | 20 | 40856 |
| MSB526 | mirEx197; syb2541 | ctrl | 6,S8 |  | 10 | 20154 |
| MSB526 | mirEx197; syb2541 | auxin | 6, S8 |  | 19 | 36135 |

**Table S2.** CRE activity reporter. Strains and their properties used as recombination reporter to study the efficiency and specificity of the cell-type specific CRE recombination.

| Strain | Promotor | Ref. | Neuron | # Animal | # cells/ Animal | # expected cells | % Animal | comments |
| --- | --- | --- | --- | --- | --- | --- | --- | --- |
| MSB205 | *rgef-1p* | [25] | PanNeuro | 20 | many | 300 | 100 | individual cells cannot be counted reliably |
| MSB211 | *unc-4p* | [25] | A-type | 10 | 10-25 | 12+9 | 100 | visible in the ventral nerve chord |
| MSB495 | *acr-5p* | this study | B-type | 24 | 18-22 | 11+7 | 100 | visible in the ventral nerve chord, recombination also visible in a few cells in the tail. |
| MSB213 | *unc-25p* | [25] | D-type | 10 | 16-19 | 13+6 | 100 | visible in the ventral nerve chord |
| MSB214 | *myo-3p* | [25] | BWM | 20 | all? | 95 | 100 | individual cells cannot be counted reliably |
| MSB282 | *flp-22Δ4p* | this study | SMD | 20 | 4-8 | 2+2 | 100 | In the majority of the animals the recombination is happening in the head neurons, SMD with a few other false positive in the other head neurons and a few animals with a false positive with 1-2 tail neurons. |
| STR335 | *des-2p* | [109] | PVD | 10 | 6 | 2+2 | 100 | additional recombination observed in FLP; m4; tail neurons |
| STR655 | *mec-17p* | [109] | TRN | 10 | 6-8 | 6 | 100 | false positive presumably due to transient expression of CRE in PVD in the mec-17 promotor |
| MSB210 | *nlp-12p* | this study | DVA | 10 | 1-2 | 1 | 100 |  |
| MSB500 | *F49H12.4p* | this study | PVD | 28 | 3 | 4 | 100 | possible recombination in AQR, but cannot be seen in these animals because of the myo-2::mCherry markers |

**Table S3.**
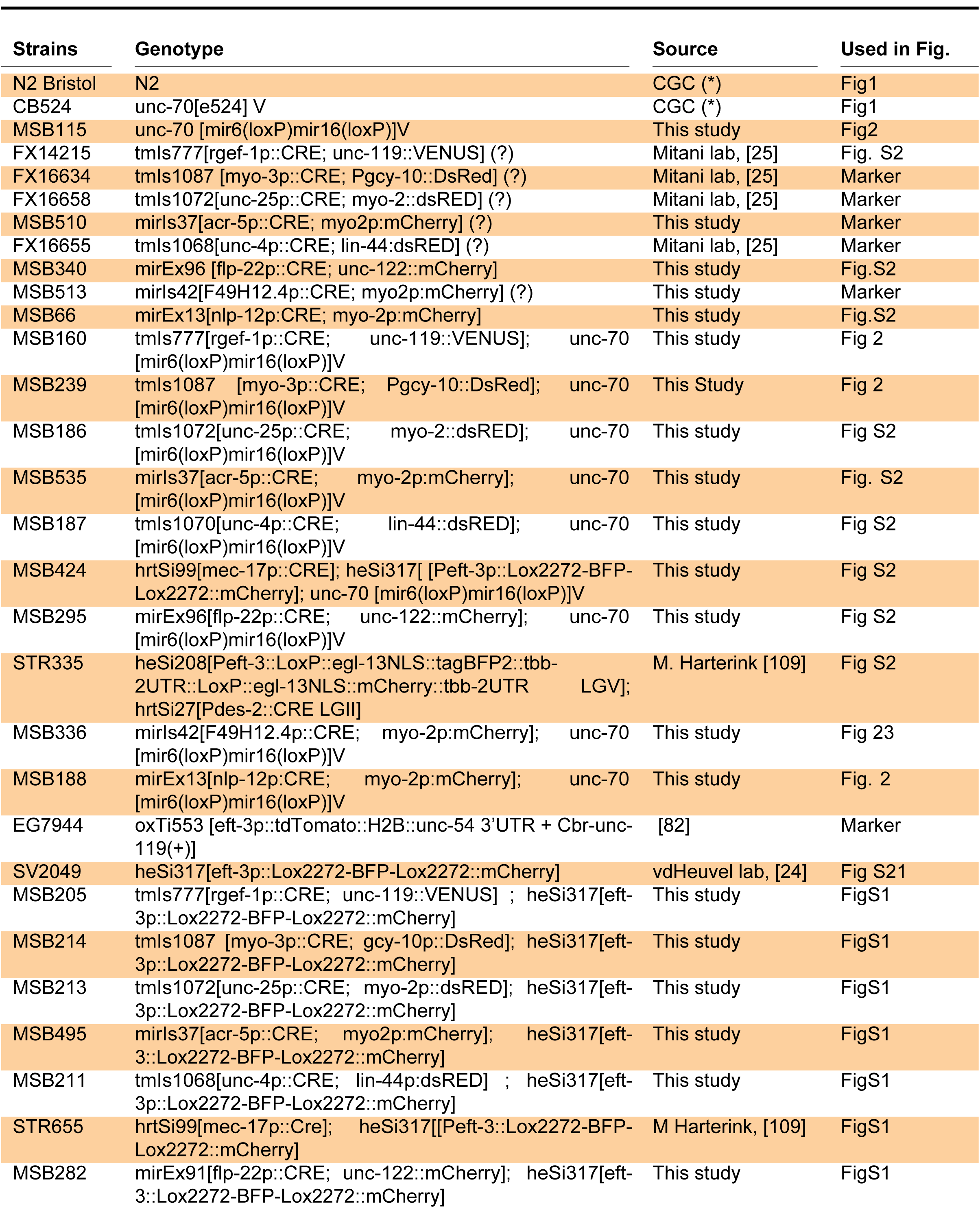
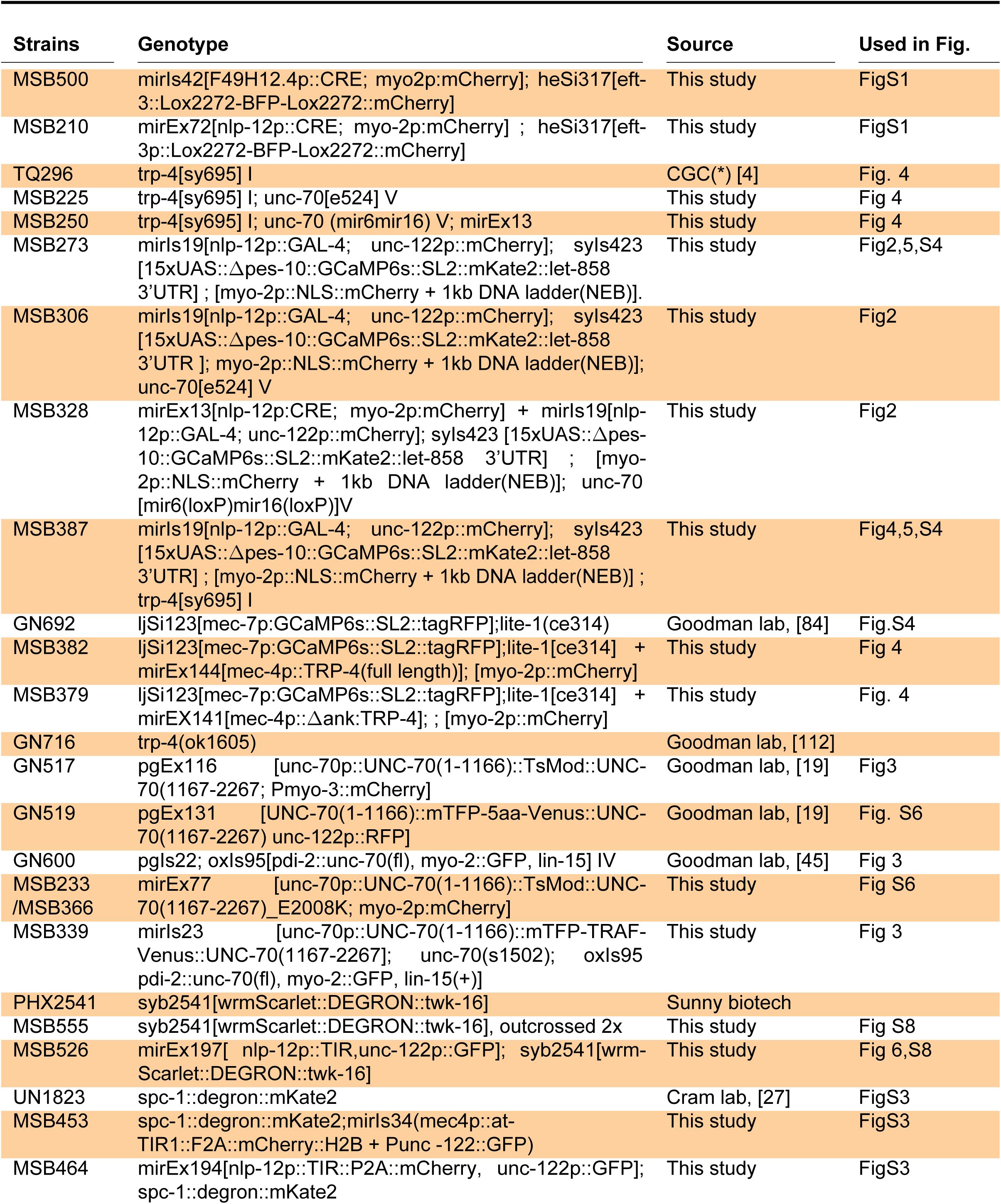
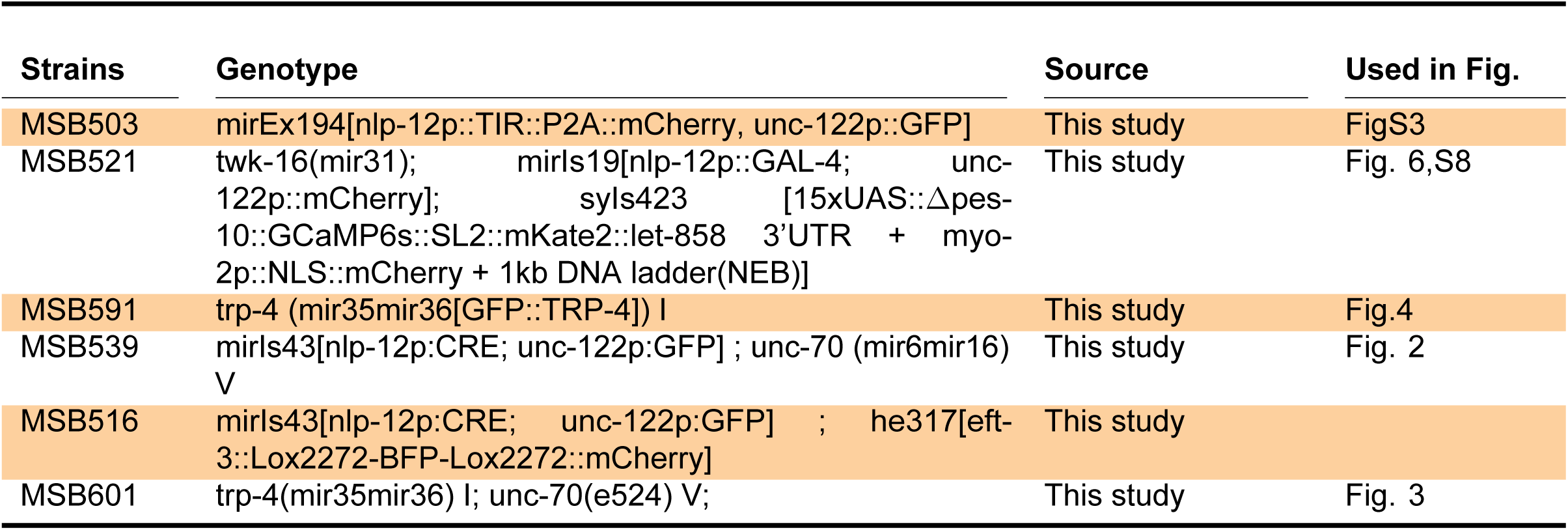
Strains used in this study.

